# The Microbiome and Metatranscriptome of a Panel from the *Sarracenia* Mapping Population Reveal Complex Assembly and Function Involving Host Influence

**DOI:** 10.1101/2024.06.10.598016

**Authors:** Jiazhang Cai, Iqra Mohsin, Willie Rogers, Mengrui Zhang, Lin Jiang, Russell Malmberg, Magdy Alabady

**Author notes:** Corresponding author: Magdy Alabady. January 2023 (Russell Malmberg).

## Abstract

*Sarracenia* provide an optimal system for deciphering the host-microbiome interactions at various levels. We analyzed the pitcher microbiomes and metatranscriptomes of the parental species, and F1 and F2 generations from the mapping population (*Sarracenia purpurea* X *Sarracenia psittacina*) utilizing high-throughput sequencing methods. This study aimed to examine the host influences on the microbiome structure and function and to identify the key microbiome traits. Our quality datasets included 8,892,553 full-length bacterial 16s rRNA gene sequences and 65,578 assembled metatranscripts with microbial protein annotations. The correlation network of the bacterial microbiome revealed the presence of 3-7 distinct community clusters, with 8 hub and 19 connector genera. The entire microbiome consisted of viruses, bacterial, archaea, and fungi. The richness and diversity of the microbiome varied among the parental species and offspring genotypes despite being under the same greenhouse environmental conditions. We have discovered certain microbial taxa that are genotype-enriched, including the community hub and connector genera. Nevertheless, there were no significant differences observed in the functional enrichment analysis of the metatranscriptomes across the different genotypes, suggesting a functional convergence of the microbiome. Our results revealed that the pitcher microcosm harbors both rhizosphere and phyllosphere microbiomes within its boundaries, resulting in a structurally diverse and functionally complex microbiome community. A total of 50,424 microbial metatranscripts were linked to plant growth-promoting microbial proteins. We show that this complex pitcher microbiome possesses various functions that contribute to plant growth promotion, such as biofertilization, bioremediation, phytohormone signaling, stress regulation, and immune response stimulation. Additionally, the pitcher microbiome exhibits traits related to microbe-microbe interactions, such as colonization of plant systems, biofilm formation, and microbial competitive exclusion. In summary, the demonstrated taxonomical divergence and functionally convergence of the pitcher microbiome are impacted by the host genetics, making it an excellent system for discovering novel beneficial microbiome traits.

## Introduction

Plants live in association with a wide range of microorganisms, including bacteria, fungi, protists, nematodes, and viruses, all referred to as the plant microbiome. In natural habitats, through this association, microorganisms play a significant role in enhancing plant growth, productivity, and health (Trivedi et al. 2020). Experimental evidence has demonstrated that plants have the ability to modify the abundance and diversity of microbiomes in their environments (Mahnert et al. 2015). Nevertheless, the precise mechanisms through which this occurs remain unidentified. Plants may have acquired genetic traits that impact their interaction with their microbiomes. The identification of these traits and their mechanisms are essential for the success of the ongoing efforts to harness and manipulate microbiomes in order to optimize their advantages for the host plants (Afridi et al. 2022; Ke et al. 2021; Morales Moreira et al. 2023).

Carnivory evolved independently on 13 occasions in flowering plants, which includes four instances in monocots (three in Poales and one in Alismatales) and nine instances in eudicots (three in Caryophyllales, three in Lamiales, two in Ericales, and one in Oxalidales). There is a total of 810 species belonging to 21 plant genera, which are distributed across 13 families and six orders (reviewed in (Fu et al. 2023)). As a result of convergent evolution, some carnivorous species have developed modified leaves that allow them to capture and kill insect prey. These species also host a complex microbiome that aids in the digestion of the captured prey and support plant growth through an array of direct and indirect traits. These traits might be considered as extensions of the host plant traits, which are equally significant to the inherent plant traits (the concept is reviewed in (Whitham et al. 2006, 2008)). The “pitfall” carnivorous plants evolved to trap insects in three separate lineages (Albert et al. 1992). Using modified leaves called ‘pitchers’, they digest insect prey with the aid of a complex microbiome that develops inside their pitcher’s fluid. In the genus *Sarracenia*, species differ in their pitcher morphology, which in turn affects the insect species they capture for nutrients (Ellison et al. 2004, 2003a), and also affects the initial assembly of the pitcher microbiomes. The factors involved in shaping the microbiome community likely involve host plant genetics, local environmental conditions, as well as prey’s microbiomes. The interactions between the *Sarracenia*, its microbiome, and captured prey likely played a role in driving the process of plant speciation. In return, the host species genetics may have evolved traits to impact the composition and function of their microbiome communities. The host impact can be measured by examining the taxonomic divergence and functional convergence of the pitcher microbiome, which are two prevalent phenomena in microbial systems (Louca et al. 2018, 2016).

The bacteria, archaea and eukaryotes that reside in pitcher have been described by organismal methods (Ellison et al. 2003b; Baiser et al. 2013) and by environmental sequencing (Adlassnig et al. 2011; Boyer and Carter 2011; Koopman and Carstens 2011). Multiple studies have explored the impact of the *Sarracenia* host plant on the microbiome assembly by comparing the microbial communities in natural and artificial pitchers (Koopman et al. 2010; Bittleston et al.; Ellison et al. 2021; Grothjan and Young 2022), as well as among various *Sarracenia* species (Heil et al. 2022). These studies concluded that *Sarracenia* may play an active role in shaping the structure of their microbiome. However, is this impact is regulated in part by the genetic makeup of the host (i.e., deterministic impact)? To address this question, we are using a *Sarracenia* mapping population in a controlled environment to characterize the microbiome diversity, structure and function among genotypes of same genetic background.

Malmberg *et al*. developed a *Sarracenia* genetic linkage map of 437 SNP and SSR markers with a total length of 2017 cM using an F2 generation of 280 plants from a genetic cross between *Sarracenia purpurea* (*Spu*) and *Sarracenia psittacina (Spa)* (Malmberg et al. 2018). They mapped 64 pitcher QTLs traits that were involved in prey attraction, capture and kill, and digestive processes/mechanisms. Despite the fact that the morphological characteristics of these traits are indicative of their adaptation to prey capture, their impact on the structure and function of the microbial community remains uncertain. We hypothesize that *Sarracenia* evolved genetic features for interacting with their vital microbiomes in a manner similar to the convergent evolution observed in their leaf structure.

Pitcher plants are ideal for studying host-microbiome and microbiome-microbiome interactions at the experimental and molecular levels for several reasons. Firstly, each fluid-containing pitcher is a natural microcosm with well-defined boundaries, making it experimentally amenable. This differs from other symbiotic microbial systems that are less amenable to experimentation, such as gut microbiomes, or those that do not have well-defined boundaries such as plant rhizosphere and phyllosphere microbiomes. Secondly, different species within the *Sarracenia* genus have varying leaf traits associated with differing microbial communities, that attract and digest different species of insects (Grothjan and Young 2019). These genetically determined characteristics may have allowed pitcher plants to impose control over the microbiome assembly in their pitchers. Thirdly, the various species of *Sarracenia* can hybridize readily, offering a unique opportunity to incorporate a genetic approach to examine plant genome influences on microbiome assembly (Furches et al. 2013). Although pitcher plants have long attracted biologists, we know little about the assembly mechanisms involved in the microbiome assembly and structure in their pitchers (Grothjan and Young 2019). It has been suggested that factors other than prey capture and colonization by eukaryotic species may affect the recruitment of bacteria to their pitchers (Grothjan and Young 2019).

The objectives of this study are to evaluate the variations in the microbiome structure and function among different genotypes of the *Sarracenia* mapping populations, namely *Spu*, *Sps*, F1 and F2, and to characterize the composition and functions of the *Sarracenia* microbiome. For a panel of these genotypes, we utilized the single molecule (Oxford Nanopore Technology, ONT) and short reads (Illumina) technologies to sequence the full-length 16S rRNA gene (microbiome) and total mRNA (metatranscriptome), respectively, from their pitcher fluids (Figure 1A). We employed a variety of specialized databases, as well as bioinformatical tools to analyze the data (Figure 1B-D).

**Figure 1:**
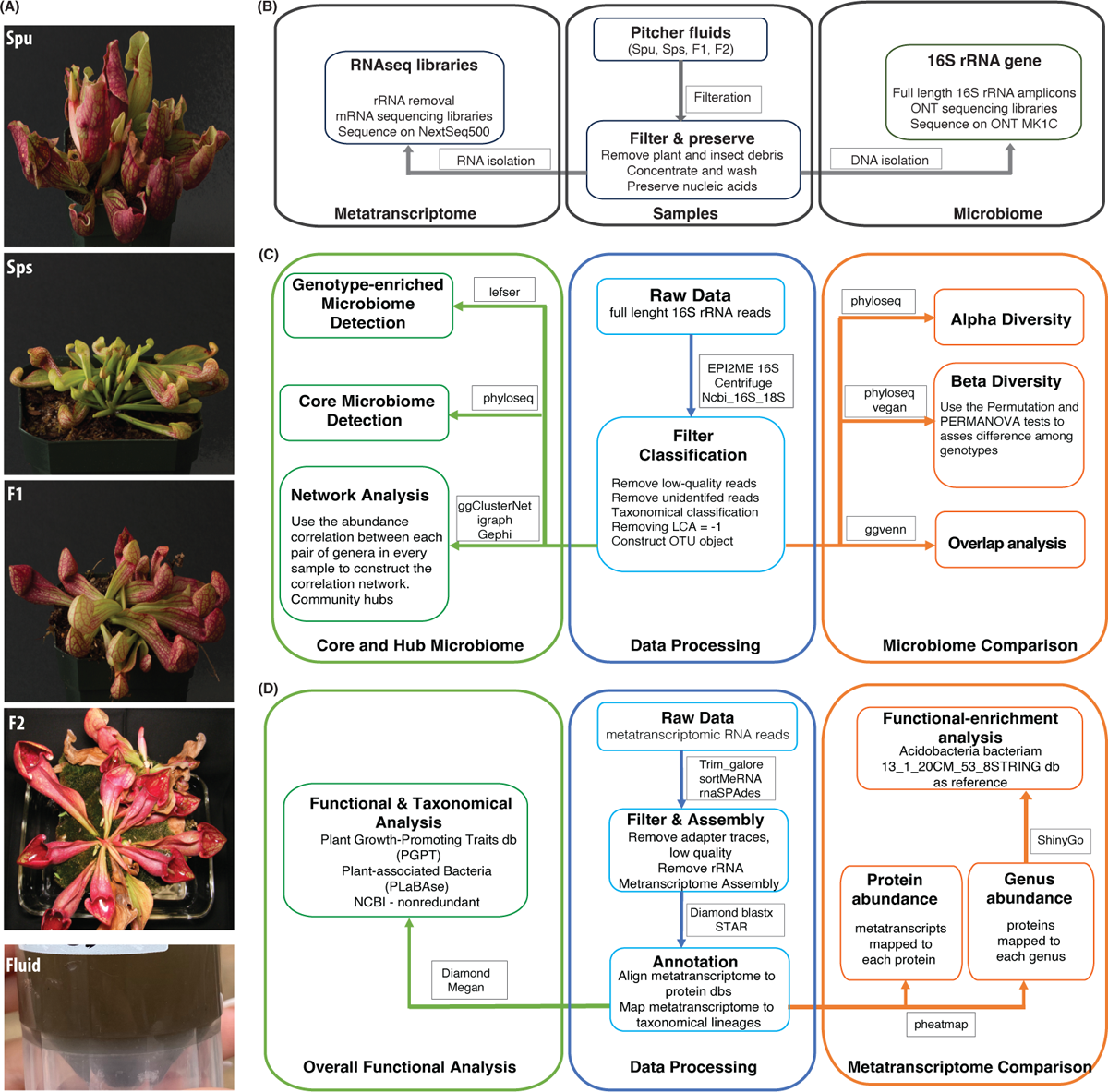
Summary of the experiment and analysis pipelines. (A) Photos of the parents (*Spu* and *Sps*) and selected F1 and F2 individuals of the *Spu X Sps* mapping popula{on, and a sample of the collected pitcher fluid. (B) The experimental process of preparing the pitcher fluid samples and sequencing. (C) Modules of the 16S rRNA microbiome analysis pipeline. (D) Modules of the metatranscriptome RNA analysis pipeline.

## Material and Methods

### Genotypes

The samples of parental species are true biological replicates because they are clones of the same individuals, but the samples from the F1 and F2 generations are separate individuals. To simplify the study, we treated the F1 individuals as biological replicates of each other and referred to them as “F1”. Similarly, the F2 individuals were treated as biological replicates and referred to them as “F2”.

### Collecting *Sarracenia* Pitcher Fluid

We collected the fluid from the pitcher leaves of *Spu*, *Sps*, F1 and F2 plants, which were grown under the same environment in the greenhouse (Figure 1A). All pitchers were at least 8 weeks old to ensure stable microbial communities. The pitcher fluid was collected using sterile glass pipettes. For *Spu*, F1 and F2 plants, the glass pipettes were easily inserted into the pitchers as the flap is open. The *Sps* plants have a narrow body and a closed flap, which required a small incision in the flap to insert the glass pipette into the pitcher (Figure 1A). Due to the morphology of *Sps*, some pitchers were dry or contained a small amount of fluid. In these cases, sterilized water was added to elute the microbiome that is attached to pitcher’s internal surface. The fluid samples were collected into 50 ml conical sterilized tubes and stored in a −20 ℃ freezer.

### Sample Preparation

We thawed the pitcher fluid samples on ice for 30-40 minutes, followed by filtration using the Steriflip Sterile Centrifuge Tube Top Filter Unit (Cat# SCNY00020) to remove any plant material and dead insect matter. A schematic diagram of the sample preparation is shown in Figure 1B. The filtered fluid was a cloudy and homogenous mixture, indicating the presence of microbial growth. We centrifuged the samples at 3000 rpm at 4℃ for 20 minutes to precipitate the microbiota, discarded the supernatant, suspended the pellet in 450 µl of PBS buffer, and mixed by pipetting. We added 750 µl of DNA/RNA Shield (Cat# R2002) to the suspension and mixed by pipetting and inverting. The material was transferred into a bead-bashing lysis tube and battered at high frequency for 2 minutes. Next, the samples were then centrifuged for 30 seconds at 16,000 rpm, divided into 400 µl aliquots in 2 ml tubes with 2 volumes of the DNA lysis buffer added to each tube. Finally, we used the ZymoBIOMICS DNA/RNA Miniprep Kit (Cat# R2002) to isolate both DNA and RNA.

### Microbiome DNA Isolation

We transferred the entire sample into a SpinAway Yellow Filter unit, followed by centrifugation at 16,000 rpm for 30 seconds. The flow-through, which contained the RNA, was saved in a 15 ml tube. An equal volume of 100% ethanol was added, and the tube was placed on ice. After processing the entire sample, a new collecting tube was used and 400 µl of DNA/RNA Prep Buffer was added to the column, followed by centrifugation at 16,000 rpm for 30 seconds, adding 700 µl of DNA/RNA wash buffer to the column, and centrifugation at 16,000 rpm for 30 seconds. Another 400 µl of the wash buffer was added to the column followed by centrifugation at 16,000 rpm for 2 minutes. We then carefully transferred the yellow filter into a nuclease-free 1.5 ml tube and added 50 µl of DNase/RNase-free water directly in the center of the yellow column matrix and incubated it at room temperature for 5 minutes. While waiting, we assembled a Zymo-Spin III-HRC Filter unit and added 600 µl of ZymoBIOMICS HRC Prep Solution into it, followed by centrifugation at 8000 rpm for 3 minutes. The elution column was then centrifuged at 16,000 rpm for 30 seconds, and the eluted DNA was transferred into the prepared HRC filter unit and centrifuged for 16,000 rpm for 3 minutes to elute the final DNA into1.5 ml tube.

### Microbiome RNA Extraction

We transferred the RNA sample into a SpinAway Green Filter unit, followed by centrifugation at 16,000 rpm for 30 seconds, replacing the collection tube, adding 400 µl of DNA/RNA buffer to the column, mixing by pipetting, and centrifugation at 16,000 rpm for 30 seconds. We added 700 µl of DNA/RNA wash buffer to the column and centrifuged at 16,000 rpm for 30 seconds, followed by adding another 400 µl of the wash buffer to the column, centrifugation at 16,000 rpm for 2 minutes. We then carefully transferred the green filter into a nuclease-free 1.5 ml tube, added 30 µl of DNase/RNase-free water directly in the center of the yellow column matrix, and incubated it at room temperature for 5 minutes. While waiting, we assembled the Zymo-Spin III-HRC unit, added 600 µl of HRC prep solution, and centrifuged at 8000 rpm for 3 minutes. The green column matrix was centrifuged at 16,000 rpm for 30 seconds to elute the RNA into the nuclease-free tube. The eluted RNA was transferred into the prepared HRC filter in another 1.5 ml nuclease-free tube and centrifuged at 16,000 rpm for 3 minutes to elute the final RNA in 1.5 ml tube. The RNA tubes were labeled and stored at −80°C.

### DNA and RNA Quality Assessment

We used the Fragment Analyzer Long Fragment (LF) assay to examine the DNA quality and the Bioanalyzer Pico RNA Assay to assess the RNA integrity. The concentrations were assessed using the Qubit 4 fluorometer high-sensitivity assays.

### Full-length 16S rRNA Gene Libraries and Sequencing

we used the Oxford Nanopore Technology (ONT) amplicon sequencing method to build the 16s rRNA gene libraries and sequence them on the MinIon Mk IC platform. First, we built the libraries using the 16S barcoding kit (SQK-16S024) for rapid sequencing amplicons following the manufacturer protocol. Briefly, the ONT primers (F: TTTCTGTTGGTGCTGATATTGCAGRGTTYGATYMTGGCTCAG, and R: ACTTGCCTGTCGCTCTATCTTCRGYTACCTTGTTACGACTT) were used to amplify the full-length rRNA gene from the isolated microbiome DNA samples. The amplicons (∼ 1500 bp) were barcoded by PCR using the ONT barcoding primers (SQK-16S024), followed by multiplexing, and ONT adapter ligation. The Final library pool was sequenced on the Mk IC platform using FLO-MIN106 flow cells. The final pool of 16S full-length ONT libraries included 13 samples (4 *Spu*, 3 *Sps*, 3 F1, and 3 F2). One of the three F2 samples failed in the sequencing, leaving only two F2 samples.

### Metatranscriptomic Library Preparation and Sequencing

We treated the metatranscriptome total RNA with the FastSelect 5S/16S/23S reagent (Qiagen, cat# 335921) to deplete the rRNA following the manufacturer protocol. We started with 12.5µl total RNA, fragmented for 2 min at 89 °C, and eluted the fragment RNA in 18 µl RNase-free water. The rRNA depleted was used immediately to prepare the metatranscriptome libraries using the SEQuoia complete RNAseq kit (Bio-Rad cat# 17005726) following the manufacturer’s procedures without any modifications. The final sequencing libraries were assessed on the Fragment analyzer (Figure S1), pooled equimolarly, and sequenced on the NextSeq 500 sequencer using a mid-output kit. The sequencing run produced over 171 million reads.

### Microbiome Data Analysis

#### Taxonomical Classification and Abundance

We carried out the primary analysis using the EPI2ME 16S workflow (ONT 16S workflow 2019) to generate the taxonomic classification and abundance of the 16S genes for all samples. The 16S workflow uses the Centrifuge algorithm (Kim et al. 2016) to map the full-length 16s reads to the NCBI 16S-18S database and classify them according to the taxonomy of their best matches. The obtained taxonomical matrix for all samples was then used for more in-depth analysis in R as described in the results section. Figure 1C illustrates the overall analysis pipeline, which is divided into three modules: data preprocessing, microbiome comparison, and core and hub microbiome analyses. The R packages and analysis tools utilized in this study pipeline are detailed in Figure 1C and will be elaborated on in the results section.

#### Alpha and Beta Diversity

We utilized the Alpha diversity test to evaluate microbial diversity within each group, that is, among replicates of the same genotype (4 Spu, 3 Sps, 3 F1, and 2 F2). The alpha diversity was assessed using three methods: Chao1 (Chao 1984), Pielou’s Evenness (Volvenko 2014), and Shannon (Shannon 2001). The pairwise Wilcoxon test (Wilcoxon 1992) was then used to test the significance of the variance of the alpha diversity among genotypes. The difference in the microbiome composition among genotypes was calculated using beta diversity analysis, followed by visualization using *PCoA* (Principal Coordinate Analysis) (Gower 1966). The sample distances were calculated using Bray-Curtis distances (Bray and Curtis 1957; Kers and Saccenti 2022).

#### Ubiquitous, Genotype-enriched, and Core Microbiomes

We used an abundance-based approach to identify ubiquitous microbiomes, which is the microbiome existing in all samples above the background noise. This was followed by an overlap analysis using the ggvenn R package (Yan 2021) to identify the genotype-enriched microbiomes. As for the core microbiome, we used the microbiome R package (Lahti and Shetty 2018) to identify the core pitcher microbiome after optimizing the prevalence and limit of detection factors. We then employed the Linear Discriminant Analysis Effect Size (LEfSe) (Segata et al. 2011) method to identify the most distinctive microbial genera showing significant differential abundance between each pair of genotypes. The LefSe method is powerful because it combines statistical significance with the evaluation of biological consistency (effect size) using several tools including the Kruskal-Wallis test (Kruskal and Wallis 1952) and Linear Discriminant Analysis (LDA) (Fisher 1936).

#### Network and hub microbiome

We implemented network analysis to gain insights into the structure of the pitcher microbiome community. We deployed ggClusterNet (Wen et al. 2022) to infer the correlation network followed by visualization using Gephi (Bastian et al. 2009). Subsequently, we computed three scores for each node: hub score, within-module connectivity Z-score, and participation coefficient scores. The scores reflect the node’s connectivity to all other nodes in the network, to nodes within the same community, and the evenness of edges distribution within each community, respectively. The network structure, and hence all three node scores, will change when the number of nodes, the threshold for correlations, and the *p-values* are altered. Therefore, we examined the changes in the network structure across various correlation criteria, *p-*values, and node counts (Figure S2). As a result, we set the correlation criterion to 0.7 and the p-value cutoff to 0.05.

### Metatranscriptome Data Analysis

#### Metatranscriptome Assembly and Annotation

The metatranscriptome analysis pipeline is shown in Figure 1D. First, we trimmed the sequencing reads to remove adapter traces as well as short and low-quality reads using Trim_galore. Then we used the rRNA database v4 and sortMeRNA (Kopylova et al. 2012) to remove rRNA reads from each sample. The remaining metatranscriptomic mRNA reads from all samples were combined and assembled using rnaSPAdes (Bushmanova et al. 2019). The assembled metatranscriptome was deduplicated to remove identical metatranscripts followed by annotation against a customized uniprot_sprot protein database, which included bacterial, fungal, viral and archaea sprot sequences (uniprot_sprot_microbial) using diamond blastx (*--top 1--evalue 0.05*). Also, we mapped the metatranscriptome to the NCBI nonredundant and GTDB protein databases using diamond blastx to obtain accurate taxonomical lineages. Subsequently, we mapped the cleaned sequence reads of each sample to the annotated metatranscriptome using STAR (Dobin et al. 2013) and compiled the count matrix for all samples with taxonomical information. Lastly, we converted the count matrix to CPM matrix.

#### Abundance of Genera and Metaproteins

In order to determine the abundance of taxa, the count matrix was transformed from reads per protein to genus abundance. The genus abundance was determined by counting all metatranscripts mapped to the proteins of the genus (according to the uniprot_sprot annotation) for the *Spu*, *Sps*, F1, and F2 samples. Next, we calculated the *z*-core and *p*-value for each genus count in each sample and filtered out all genera with *p*-values > 0.05. The TMM method was used to normalize the abundances of the filtered genera. Similarly, in order to identify prevalent proteins, the metatranscripts count matrix was transformed into a metaprotein abundance matrix. The metaprotein abundance was determined by counting the metatranscripts mapped to the same protein, based on the uniprot_sprot annotation, from multiple genera. Next, we calculated the *z*-core and *p*-value for each metaprotein count in each sample and filtered out all metaproteins with *p*-values > 0.01, followed by a TMM normalization.

#### Functional Core Microbiome

We used two criteria to identify the functional core microbiome: prevalence > 0.4 and limit of detection > 50 metatranscripts. Unlike the structural core microbiome, which was identified based on the 16S rRNA gene data, the functional core microbiome relies on the prevalence and abundance of protein-coding metatranscripts, hence their functions.

#### Genotype Functional-enrichment Analysis

We conducted function category enrichment analysis on the proteins that were enriched in each genotype (*p*-value < 0.05) using ShinyGO 0.80 (Ge et al. 2020) and *Acidobacteria Bacteriam* 13_1_20CM_53_8STRINGdb as a reference. The pathway databases of the enrichment analysis included GO biological processes, GO cellular components, GO molecular functions, annotated UniProt keywords, and local network clusters (STRING). We then manually filtered the list of enriched functions to remove redundancy, which existed as a result of using multiple databases.

#### Overall Microbiome Functional Analysis

We used the intensively curated Plant Growth-Promoting Traits (PGPT) (Patz et al. 2024) from the Plant-associated Bacteria (PLaBAse) project (Patz et al. 2021) to gain insights into the major functions of the pitcher microbiome community. The metatranscriptome was mapped to the mgPGPT protein database using the Diamond blastx (mini. E-value ≤ 0.05) and used the Megan software (Gautam et al. 2023) to further analyze and visualize the results.

## Results

### Microbiome Sequencing Data

The ONT sequencing run produced 9,772,540 full-length 16S rRNA gene sequences of which 9,742,538 reads were taxonomically classified (∼ 99%) with an average classification accuracy of 93%. The read length average and mode are 1,244bp, and 1,780bp, respectively. The quality score average and mode are 13.91, and 14.05 respectively. Reads without valid barcodes were removed leaving a total of 8,999,370 sequences. The EPI2ME 16S workflow assigned a lowest common ancestor (LCA) score (Munch et al. 2008) to each full-length 16S read (File S1). The LCA score is used instead of the classical “num_genus_taxid” and is used to store records of the classification status. It has three values: 1 for a species-level classification without accurate genus classification; 0 for both species and genus classification; −1 for unsuccessful species classification. We removed 106,817 reads that had a −1 LCA score. The remaining 8,892,553 reads (minimum classification accuracy is 77%) were used in the subsequent analyses.

### Microbiome Diversity and Intergenotype Variance

The use of full-length 16S rRNA genes in this analysis offers a heightened degree of precise classification analysis, hence eliminating the necessity for data rarefaction. The non-rarefied count matrix was further normalized using the Counts Per Million (CPM) and was utilized to create the phyloseq (McMurdie and Holmes 2013) object for further examination in the subsequent analysis.

#### Alpha Diversity

As illustrated in Figure 2A, the alpha diversity is measured using Chao1 (Chao 1984), Pielou’s Evenness (Volvenko 2014), and Shannon (Shannon 2001) methods. Each method measures particular aspects of microbial diversity. Chao1 assesses the species richness, Pielou’s Evenness focuses on the evenness of species distribution, and Shannon measures both richness and evenness. All three measures reveal noticeable variations in microbial richness and evenness between the genotypes (intergenotype) and among the replicates of the same group (intragenotype). When both richness and evenness are assessed simultaneously using Shannon’s approach, the median alpha diversities of all samples are more closely aligned. The observed variations may be attributed to the limited number of replicates, variations in sequence depth among samples, genetic differences among the replicates of the F1 and F2 samples, as well as the differences in microbial diversity. We then used the pairwise Wilcoxon method (Wilcoxon 1992) to test if the alpha diversity among the parents, F1 and F2 is significantly different. All of the *p*-values are very close to 1 (File S1), indicating that none of the pairs’ diversity showed a significant difference. Furthermore, this suggests that the low replication and sequencing depth variation did not result in any statistically significant difference in the diversity and species evenness observed across the samples. In the subsequent analysis, we assume there is no significant difference in the alpha diversity among the parents, F1, and F2 generations.

**Figure 2:**
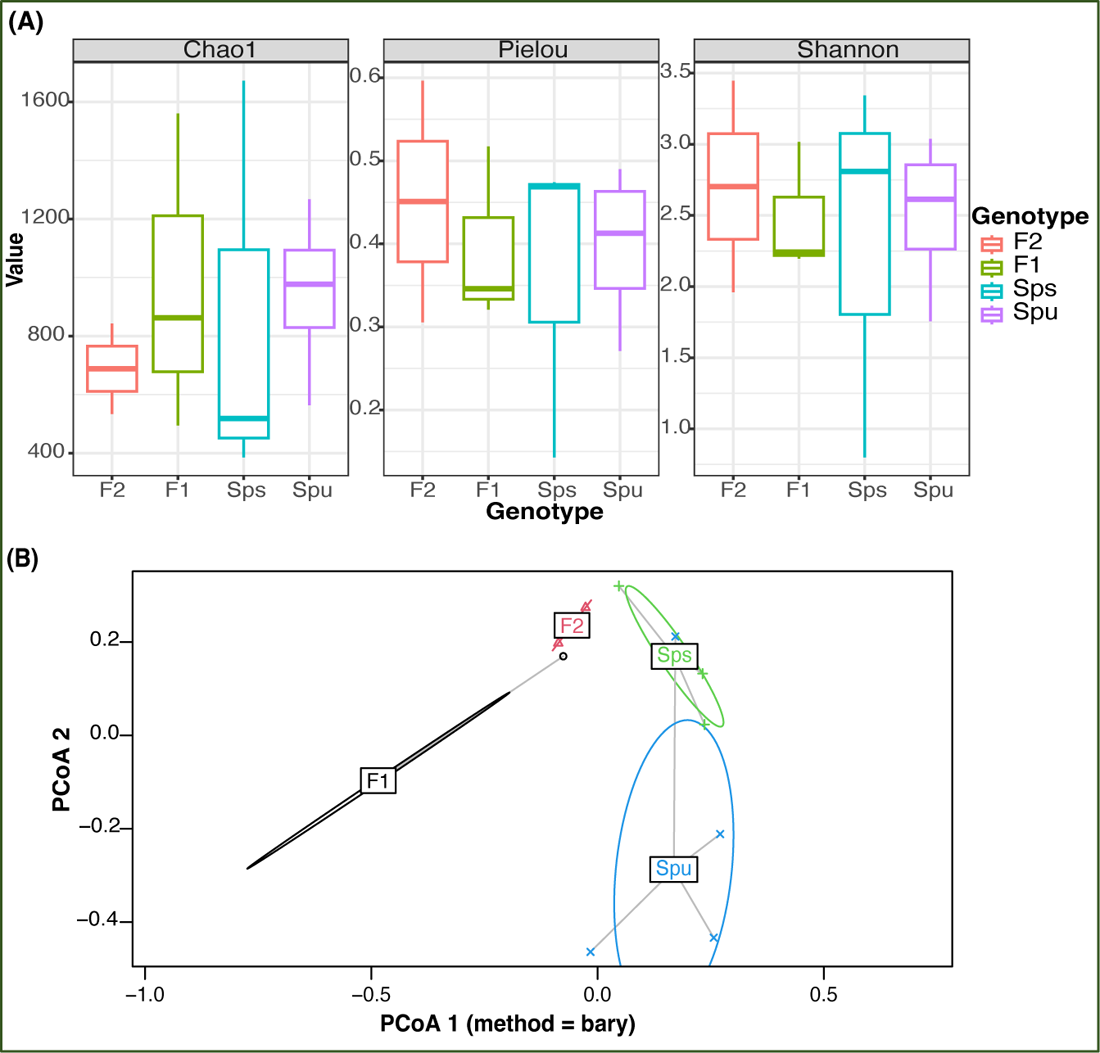
Microbiome diversity assessment. (A) Comparison of alpha diversity of different genotypes using Chao, Pielou, and Shannon tests to assess richness, evenness, and both, respec{vely. (B) Visualiza{on of the rela{onship between every sample using Principal Coordinate Analysis. The color represents the genotype of each sample. The ellipses mark the varia{on of each genotype.

#### Beta Diversity

In Figure 2B, we illustrate the beta diversity among genotypes using the *PCoA* (Principle Coordinate Analysis) (Gower 1966), where the colored ellipse depicts the dispersion of all samples from each genotype, encompassing 95% of the diversity, assuming a normal distribution. Since there are only two replicates in the F2, it is infeasible to construct an ellipse for it, and hence it uses s straight line to indicate the dispersion. The results show that the microbiome compositions of the two parent species, *Spu* and *Sps*, are different from the descendant generations, F1 and F2. To further explore the inter-genotype microbiome differences, we implemented *PERMANOVA* (Anderson 2001) to test the difference in the distribution between genotypes. The *p-*value for the test is 0.011, which indicates significant differences in microbial community structure between genotypes. We also implemented the PERMDISP test (Anderson 2006) to assess inter-genotypes microbiome dispersion. The *p-*value of the test is 0.825, indicating the microbiome dispersion is not significantly different between genotypes. Combining the results from the two tests, we can conclude that there is a significant difference in the microbial community structure between the parental species and their F1, and F2 genotypes.

### The Genotype-enriched and Core Microbiomes

We identified the genotype-enriched and core microbiomes in the pitcher fluids of the parents, F1, and F2 groups. Specifically, a genus is deemed genotype-enriched only if it is represented in every replicate of that genotype by at least one full-length 16S rRNA read. Accordingly, out of 1,960 identified bacterial genera, the pitcher microbiomes of *Spu*, *Sps*, F1, and F2 contained 180, 177, 213, and 155 bacterial genera, respectively (Table 1, File S1). There is a total of 342 non-redundant bacterial genera constituting both the ubiquitous (62), and genotype-specific (145) bacterial taxa, as well as taxa existed in two to three genotypes (135). We utilized the ggvenn R package (Yan 2021) to identify the microbiome overlap among genotypes (Figure 3A and 3C). Specifically, 62 (18.1%) bacterial genera exist in all samples of *Spu*, *Sps*, F1, and F2, among which the top 5 abundant genera are *Aeromonas*, *Rhodopseudomonas*, *Achromobacter*, *Paraburkholderia*, and *Azospirillum*.

**Figure 3:**
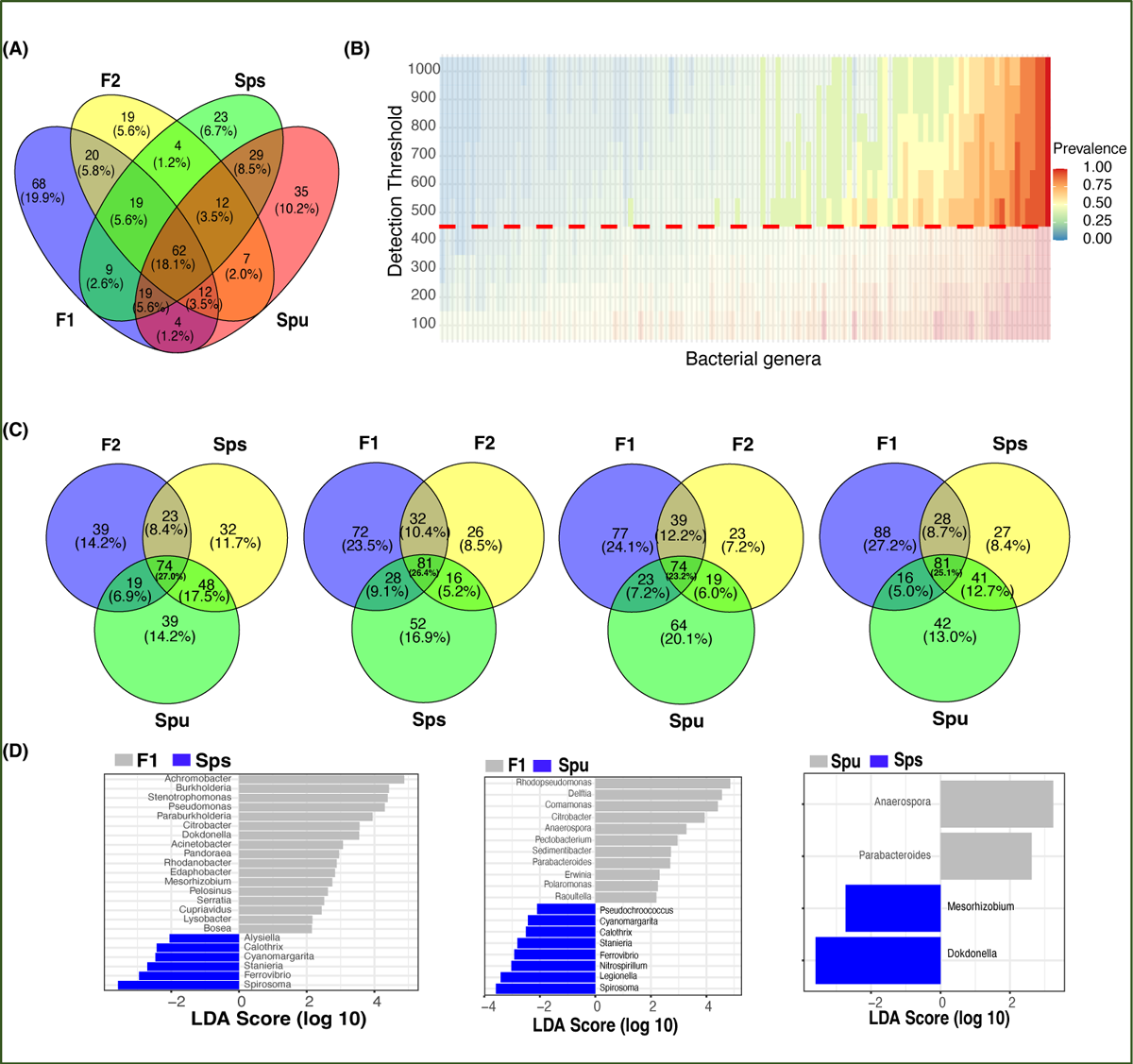
The core and genotype enriched microbiome analysis. (A) A Venn diagram showing the number of overlapped genera among *Spu*, *Sps*, F1, and F2. (B) Heatmap of the prevalence of different microbiomes under different detec{on thresholds. The detec{on value and prevalence cut-off limits are represented by the horizontal red line and transparent area, respec{vely. (C) Four Venn diagrams comparing the overlapping genera among every three genotypes to highlight the genotype specific numbers. (D) Genotype-enriched genes detected by Linear discriminant analysis Effect Size.

**Table 1:** summary of the results.

| **Bacterial 16s rRNA gene reads classification and analysis** |  |  |
| --- | --- | --- |
| Full-length classified 16S rRNA gene |  | 8,892,553 reads |
| Classified bacteria |  | 1960 genera |
| Ubiquitous and genotype-enriched (non-redundant): |  | 432 genera |
|  | *Spu* | 180 genera |
|  | *Sps* | 177 genera |
|  | F1 | 213 genera |
|  | F2 | 155 genera |
| Structural core microbiome |  | 58 genera |
| Community clusters |  | 3-7 clusters |
| Community hubs |  | 4 genera |
| Community Connectors |  | 23 genera |
| **Microbial Metatranscriptome classification and functions** |  |  |
| Metatranscripts with microbial protein annotation |  | 65,578 metatranscripts |
| Microbial genera based on metatranscriptome: |  | 2157 |
|  | Viral | 121 genera |
|  | Bacterial | 1067 genera |
|  | Archaeal | 11 genera |
|  | Fungal | 270 genera |
| Functional core microbiome |  | 90 genera |
|  | Bacterial | 68 genera |
|  | Fungal | 22 genera |
| Genotype-enriched microbes (p-value < 0.05) |  | 25 genera |
| Non-redundant microbial protein matches |  | 9,693 proteins |
| Genotype-enriched microbial proteins (p-value < 0.01) |  | 212 proteins |
| Metatranscripts mapped to PGPR proteins: |  | 50424 metatranscripts |
|  | Direct effect traits | 13,519 metatranscripts |
|  | Indirect effect traits | 29,029 metatranscripts |

For the core microbiome, we considered a genus to be a member of the core microbiome if its least prevalence under different detection thresholds is greater than 40%, as described in the microbiome analysis R package (Lahti and Shetty 2018). Unlike the ubiquitous microbiome, the core microbiome requires a specific abundance (the detection threshold) in a specific proportion of samples (prevalence) instead of in every sample. The heatmap in Figure 3B illustrates the prevalence of the core microbiome under different detection thresholds (100 to 1,000). With a prevalence equal to 0.4 and a detection limit equal to 500, a total of 58 bacterial genera constituted the core microbiome of *Spu*, *Sps*, F1, and F2 (File S1). The top 5 genera with the highest CPM normalized count in the core microbiome are *Azospirillum*, *Achromobacter*, *Rhodopseudomonas*, *Mucilaginibacter*, and *Aeromonas*. There are 42 genera that are shared by the ubiquitous and the core microbiomes, representing 67.7% and 72.4% of each, respectively (File S1).

In the genotype-enriched microbiome, 35 genera (10.2%) are unique to the *Spu* genotype, 23 genera (6.7%) to the *Sps* genotype, 68 genera (19.9%) to the F1 genotype, and 19 genera (5.6%) to the F2 genotype (Figure 3A). The results show that the microbiome of the F1 genotype differs significantly from that of the other genotypes. A similar observation can be obtained from the comparisons presented in Figure 3C. The microbiomes of the two parents (*Sps* and *Spu*) exhibit a greater similarity to each other than to either the F1 or F2 microbiomes (Figure 3A). The proportion of shared genera between *Sps* and *Spu* microbiomes is 37.8% when compared to F1 and 44.5% when compared to F2 microbiomes, which is much higher than the overlap between F1 and F2 microbiomes (Figure 3C). The overlaps between F1 and F2 microbiomes are 35.4 relative to the *Spu* and 36.8 relative to the *Sps* microbiomes. The results confirm that the *Sps* microbiome is similar to that of *Spu*, while the F1 microbiome is more closely related to the F2 microbiome (Figure 2B).

We employed LEfSe analysis (Segata et al. 2011) to examine the microbiomes of *Spu*, *Sps*, F1, and F2 to identify the most distinctive microbial genera showing significant differences in abundance across all genotype. Figure 3D displays the LEfSe analysis results for the comparisons *Sps* vs. F1, *Spu* vs. F1, and *Spu* vs. *Sps*. The F2 microbiome was excluded from the analysis due to having only two replicates, as previously explained. At a log10 LDA score of 2, there are more distinct genera in the microbiomes of F1 compared to *Spu* and *Sps*, than between *Spu* and *Sps* (Figure 3C). These results are consistent with the PCoA analysis (Figure 2B) and microbiome overlapping analysis (Figure 3A, C).

### The Microbiome Communities Networks and Hubs

This analysis included 582 genera (out of 1,960) with an accumulative normalized abundance of more than 100 across all samples. In the correlational network (Figure 4A), nodes with correlation ≥ 0.7 and *p*-value ≤ 0.05 are connected with edges. The greedy algorithm (Clauset et al. 2004) assigned the network nodes to 5 microbiome communities (clusters), which are indicated by the node and edge colors (Figure 4A-left). The edge colors represent the correlation values of connected nodes, where blue and red edges signify positive and negative correlations, respectively. Two communities only contain less than 10 genera (Cluster 1, 2), and the majority of the genera (97%) are assigned to the other three communities (Cluster 3, 4, 5). Cluster 1 (dark green) is located in the negatively correlated part of the network and serves as the hub node for the other three major microbiome communities. Cluster 2 (orange) has very few connections to the rest of the network. Cluster 3 (pink), Cluster 4 (light green), and Cluster 5 (blue) construct the main part of the microbiome network. The top 5 genera with the highest normalized count in these five communities are: *Sphingobacterium*, *Pedobacter*, *Tissierella*, *Epilithonimonas*, and *Diaphorobacter* for Cluster 1; *Gracilibacter*, *Desulfonatronobacter*, *Desulfococcus*, *Maritalea*, and *Desulfoconvexum* for Cluster 2; *Azospirillum*, *Mucilaginibacter*, *Legionella*, *Terriglobus*, and *Methylacidimicrobium* for Cluster 3; *Achromobacter*, *Rhodopseudomonas*, *Aeromonas*, *Comamonas*, and *Burkholderia* for Cluster 4; *Sphingomonas*, *Luteibacter*, *Granulicella*, *Reyranella*, and *Terrimonas* for Cluster 5.

**Figure 4:**
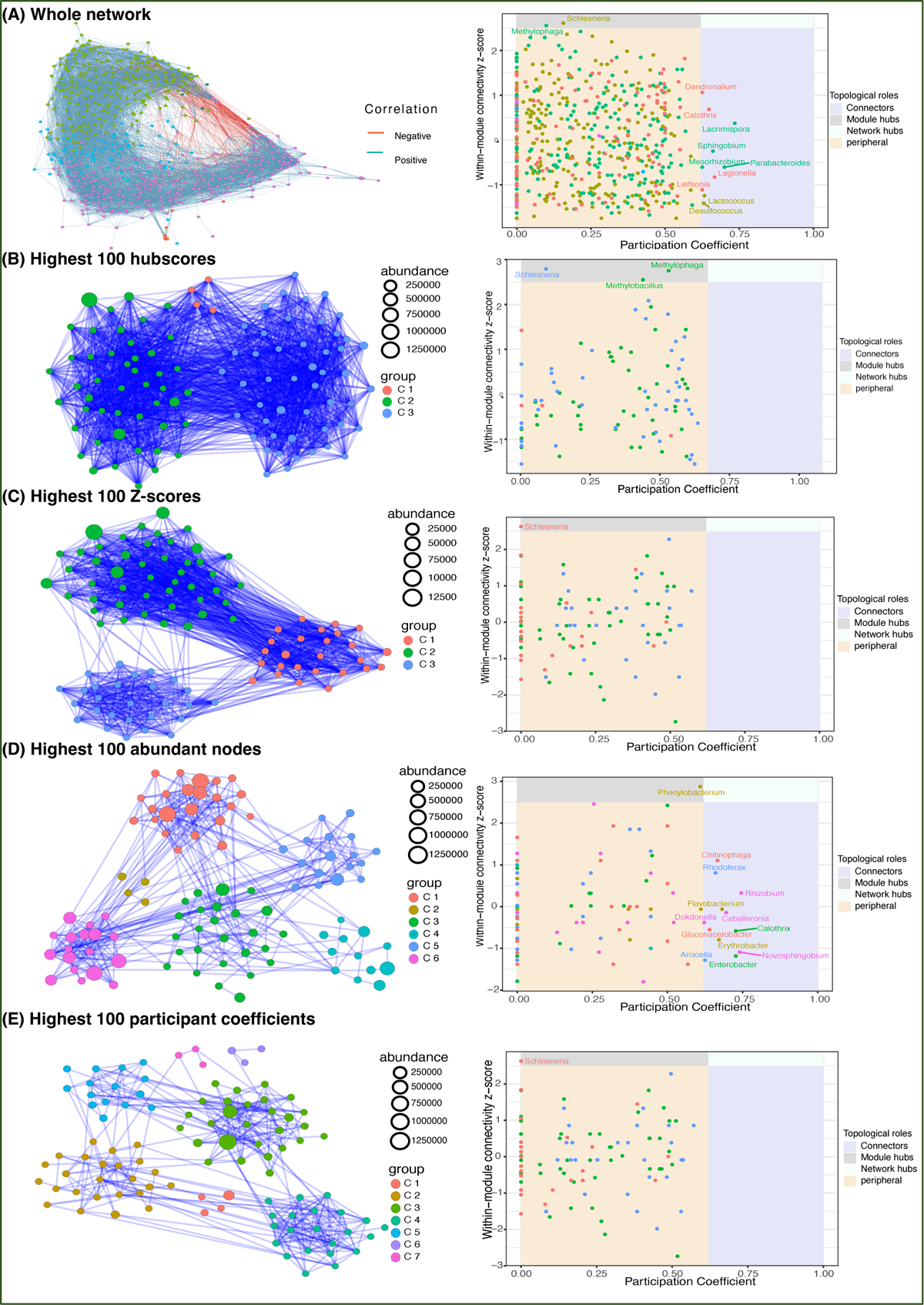
Network analysis. (A) the whole network of 582 filtered genera (leC) and connectivity scores (Right). (B-E leC) the subnetworks of top 100 genera ranked by their hubscore (B), z-score (C), normalized abundance (D), and participant coefficient (E). The connectivity scores of the four subnetworks are in their corresponding right panels.

We also computed the within-model connectivity *z*-score and participation coefficient (Guimera and Nunes Amaral 2005) for each node in the whole network, which indicates the connectivity within and between communities, respectively (Figure 4A-right). Nodes are classified into four categories based on their scores: peripheral, connections, module hubs, and network hubs (Guimera and Nunes Amaral 2005). A network hub node has a stronger connection to the network than the ordinary node. Connector and module hub nodes have more connectedness across and within communities than the average of community nodes. The remaining genera are classified as peripheral. As shown in Figure 4A-right, the whole network has 2 module hubs and 10 connectors. The module hubs are *Schlesneria* and *Methylophaga*, whereas the connectors are *Dendronalium*, *Calothris*, *Lacrimispora*, *Sphingobium*, *Mesorhizbobium*, *Parabacteroides*, *Legionella*, *Leifsonia*, *Lactococcus*, and *Desulfococcus*.

To gain a closer insight into the microbiome communities, we zoomed in on the top 100 genera ranked by their hubscore, within-model connectivity *z*-score, normalized abundance, and participation coefficient (Figure 4B-E). In the co-occurrence subnetworks (Figure 4B-E, left), node colors denote the greedy algorithm-detected clusters and node sizes represent the accumulated abundance in all 12 samples. The connectivity scores of the four subnetworks are illustrated in Figures 4B-E (right panel), respectively. The inferred networks of the top 100 genera ranked by hubscore and within-model *z*-score have similar topology including 3 highly connected community clusters each, two module hubs (*Schlesneria* and *Methylophaga*), and no connectors or network hubs (Figure 5B-C). The high connectivity in these two subnetworks can be explained by the fact that both the hubscore and within-model connectivity *z*-score prioritize the connections of each node, resulting in a highly interconnected network with fewer distinct clustering patterns. Similarly, the inferred subnetworks of the top 100 genera ranked by normalized abundance and participation coefficient (Figure 4D-E) have similar structures with 6 and 7 clusters, and *Phenylobacterium* and *Schlesneria* as module hubs, respectively. The abundance-based subnetwork has the following 11 connector nodes: *Phenylobacterium*, *Schlesneria*, *Chitinophaga*, *Rhodoferax*, *Rhizobium*, *Flavobacterium*, *Dokdonella*, *Caballeronia*, *Gluconacetobacter*, *Calothrix*, *Erythobacter*, *Novosphingobium*, *Arcicella*, and *Entrobacter* (Figure 4E-right). The abundance and participation coefficient subnetworks seem to capture more of the network topological information and identify more community connectors and hubs.

**Figure 5:**
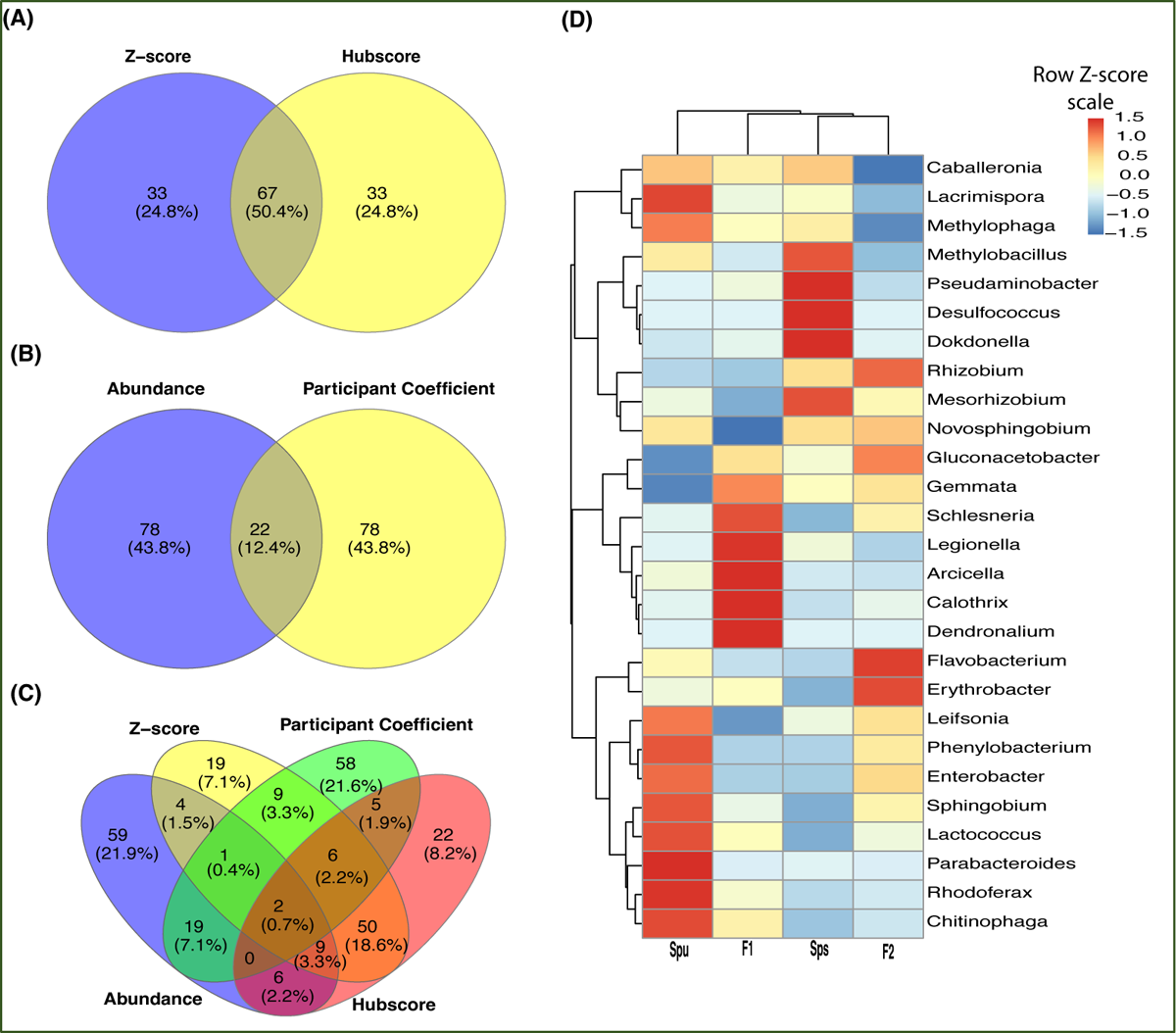
Genera overlapping among subnetworks and abundance of hub genera. (A-C) Venn diagrams showing the number of overlapped nodes (genera) among the four subnetworks. Overlapping between the hubscore and z-score top most 100 genera (A), normalized abundance and participant coefficients (B), and all four subnetworks (C). (D) A heatmap showing the normalized abundance of all hub and connector genera in the *Spu*, *Sps*, F1, and F2 samples.

The overlap among the four subnetworks is presented in Figure 5. A total of 67 genera (50.4%), including a module hub (*Schlesneria*), appeared in both hubscore and *z*-score subnetworks (Figure 5A). On the other hand, only 22 genera (12.4%) are common to both the abundance and participation coefficient subnetworks (Figure 5B). Interestingly, no module hubs or connectors were shared between these two subnetworks. Only two genera overlapped among the four subnetworks (Figure 5C). The participation coefficient quantifies the overall connectivity of each node, making it a valuable tool for capturing the structure of the network of the top 100 genera. While there is minimal overlap between the genera in the abundance and participation coefficient subnetworks (Figure 5B), both networks have a similar topology, suggesting that these two parameters represent comparable community clusters. The heatmap in Figure 5D illustrates the TMM normalized abundance of the discovered hubs and connector genera in the parents, F1, and F2 data. It reveals three clusters and six subclusters. Also, it reveals the presence of a cluster of extremely prevalent hub and connector nodes in each of the genotypes, indicating that the genotype of the host may impact the formation of the hubs and connectors of the microbiome.

### Characterization of the Pitcher Metatranscriptome

The assembled pitcher metatranscriptome included a total of 132,710 metatranscripts, with an N50, average and maximum length of 211bp, 219bp, and 8048bp, respectively. About 65,578 (49.4%), 62,382 (47%), and 63,105 (47.6%) metatranscripts have blastx protein hits (e-value ≤ 0.05) in the uniprot_sprot_microbial, NCBI nonredundant, and GTDB protein databases, respectively. Figure 6A shows a random example of the blastx alignment of metatranscripts to reference protein (min. coverage ≥ 70%). The total number of non-redundant proteins is 9693. These best protein hits come from 2157 microbial species representing 121 viral, 1067 bacterial, 11 archaeal and 270 fungal lineages (Table 1, File S2). At the microbial level, the highest represented viral clade is Riboviria (71 metatranscripts), the highest bacterial taxon is proteobacteria (17,517 metatranscripts), and an unclassified Archaea (3975 metatranscripts). About 9021 metatranscripts mapped to proteins from eukaryotic organisms, the highest among them were *Cryptophyceae* (432), *Discoba* (302), *Opisthokonta* (503), and *Sar* (4765) (Figure 6B). The metatranscriptome data revealed a functional core microbiome consisting of 90 microbial species, including 68 bacterial taxa and 22 fungal taxa (File S2). Interestingly, sixteen bacterial taxa are part of both the structural and functional microbiomes (File S2). The taxonomy tree generated from the metatranscripts (Figure 6B) offers a visual representation of the general classification of the microbial taxa in the pitcher microbiome. It is worth noting that there can be heterogeneity in the microbiome structure from one individual to another. However, as a whole, the pitcher microbiome appears to have a very intricate structure with multiple levels of interactions.

**Figure 6:**
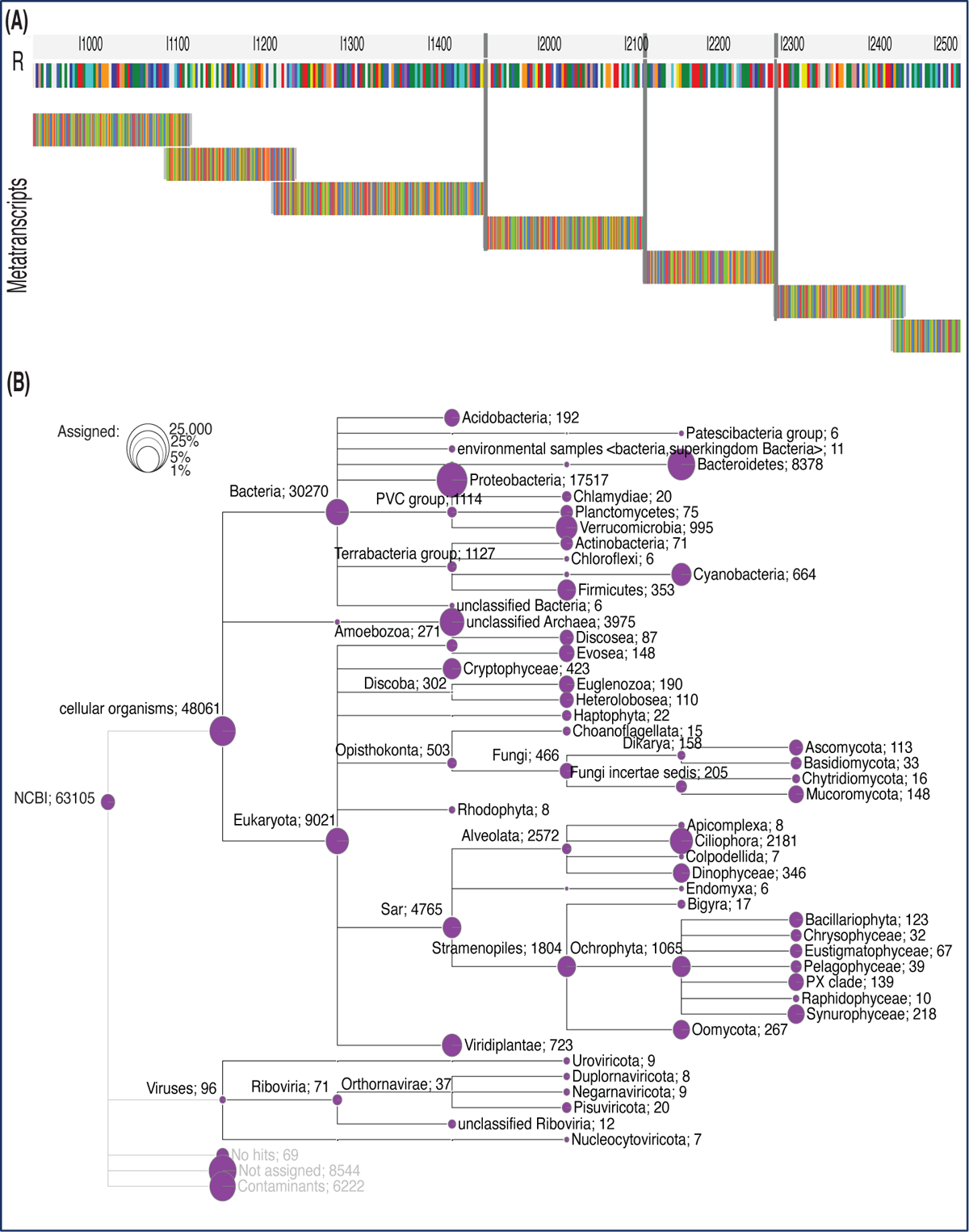
Taxonomy tree of the identified taxa based on the pitcher metatranscriptome alignment to proteins. (A) Metatranscripts of variable lengths aligned to the protein MAG TPA: carbamoyl-phosphate synthase large subunit (*Sphingobacterium* sp.). The vertical gray lines represent gaps. (B) The taxonomy tree of the identified taxa. The circles represent the log scale of number of metatranscripts assigned to the tree nodes and leaves. The numbers listed after each taxon name is the number of metatranscripts mapped to the proteins of this taxon.

### Genotype-based Differences in the Pitcher Metatranscriptomes

We examined the differences in the pitcher metatranscriptome among the genotypes (*Spu*, *Sps*, F1, and F2) using the genera and metaproteins enrichment analysis. For each genotype, we selected the genera with *p-*value < 0.05 and metaproteins with *p-*values < 0.01. There were 25 genera and 212 metaproteins meeting these criteria. The normalized abundance of these genera and proteins is illustrated in the heatmaps in Figure 7. It is evident from the heatmap clustering that there are differences in the genera abundance (Figure 7A) and metaprotein accumulation (Figure 7B) among the genotypes. Note that in Figure 7B, the heatmap includes only 25 proteins, but the complete list of proteins is provided in File S2. Despite the intergenotypic difference in the genera diversity and metaproteins, the enriched functional categories in all genotypes seem to be very similar, with small variations in the fold changes and number of genes in each category (Figure 8). For each of genotype, proteins with *p*-values < 0.01 (see materials and methods) were used in the functional category enrichment analysis. These sets included 199, 237, 236, and 201 proteins in *Spu*, *Sps*, F1, and F2 respectively. The analysis revealed a significant (*FDR* < 0.01) enrichment (more than two folds) of multiple functional categories in all genotypes. Nevertheless, there were no discernible differences in the enriched categories across the different genotypes (Figure 8), suggesting a microbial function convergence in the pitcher microcosm despite variations in structure and diversity.

**Figure 7:**
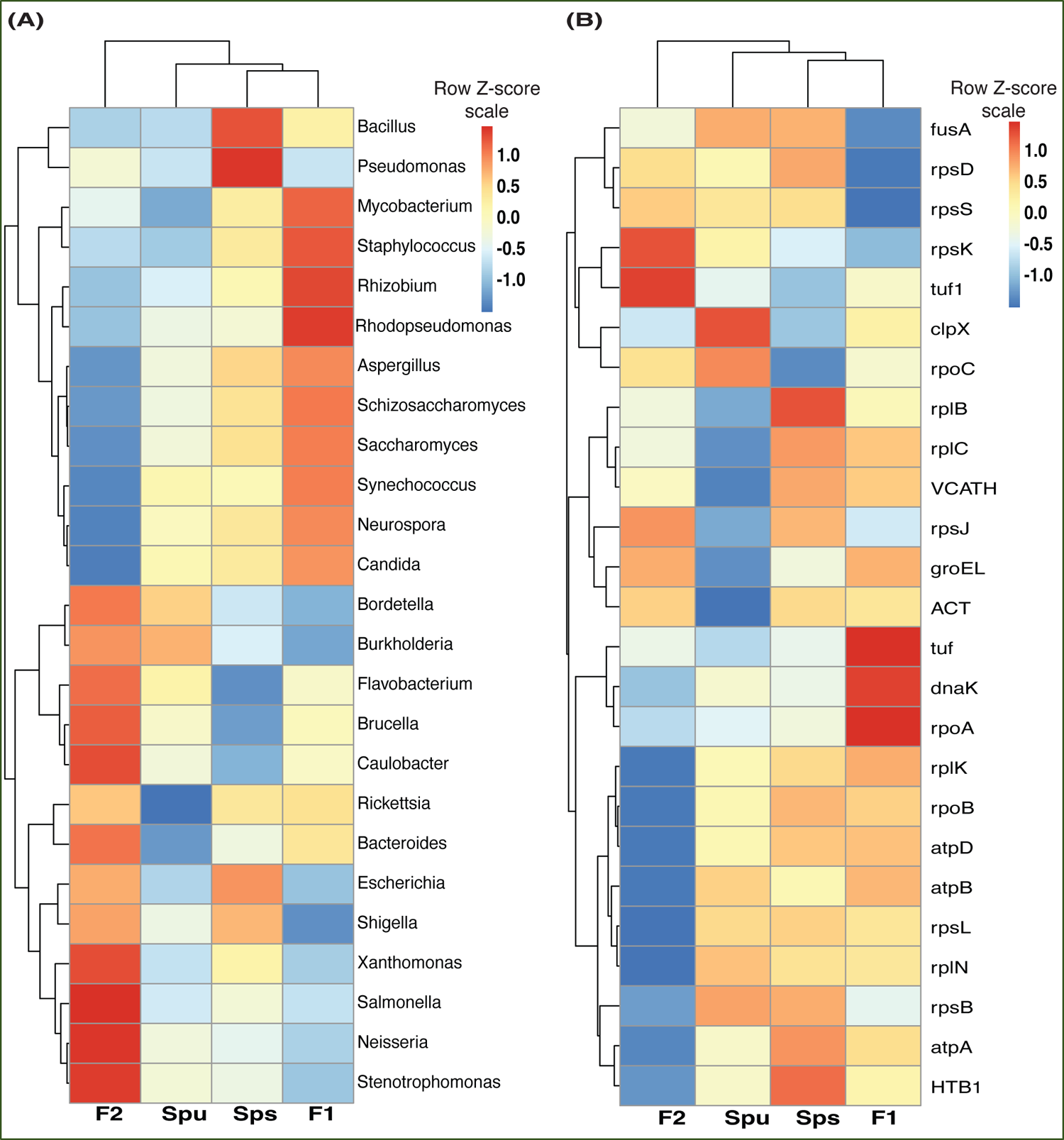
The pitcher metatranscriptome: the abundant genera and proteins showing species-related patterns. (A) heatmap showing the TMM normalized abundance of the genera in the samples of *Spu*, *Sps*, F1, and F2 (p-values < 0.05). (B) similar to “A” but showing the abundance of the 25 protein with lowest *P*-values.

**Figure 8:**
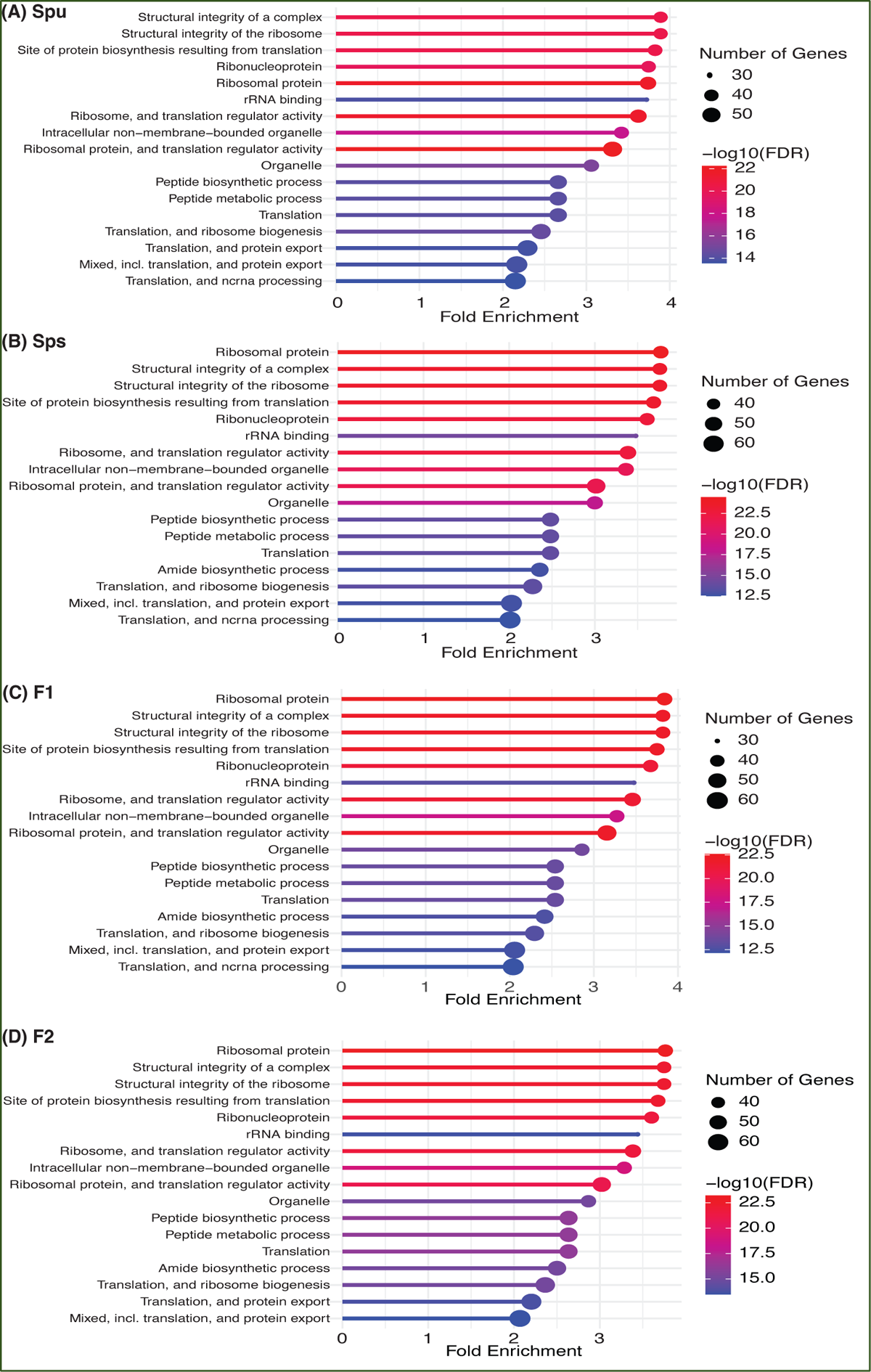
Function enrichment of the genotype-enriched metaproteins against the Go functional categories, uniprot keywords, STRING functional lists of *Acidobacteria*. The number of genes, FDR and fold enrichment of the significantly enriched functional categories in *Spu*, *Sps*, F1, and F2 metatranscriptomes are shown in A, B, C, and D, respectively.

### Phyllosphere and Rhizosphere Microbiomes

We compared the identified pitcher microbial taxa with previously reported phyllosphere and rhizosphere microbiomes (reviewed in (Bashir et al. 2022; Saeed et al. 2021)). As presented in Table 2, groups of 30, 23, and 20 microbial taxa that were reported to be performing different functions in the phyllosphere, rhizosphere, and both, respectively, were also present in the pitcher microbiomes in this study. Furthermore, many of these microbes were community connector hubs, indicating that they play a major role in microbiome assembly and functions (Table 2). The richness of the pitcher microbiome with both phyllosphere and rhizosphere microbes indicates its intricate nature and suggests that its functions extend beyond the traditional role of supporting plant nutrition.

**Table 2:** The phyllosphere (PS) and Rhizosphere (RS) bacterial (B) and fungal (F) taxa in the pitcher microbiome and metatranscriptome. The listed microbial taxa have been found in the phyllosphere and/or rhizosphere of many plant species. Here we report the coexistence of both PS and RS taxa in the *Sarracenia* pitcher fluids. D: detected, Co: community connector, and C: core genera.

| Literature<sup>1</sup> |  |  | This study |  |  |
| --- | --- | --- | --- | --- | --- |
| Microbes | Plant space | Type | Microbiome | Metatranscriptome | Core/Hub |
| *Aureobasidium* | PS | F |  | D |  |
| *Azorhizobium* | PS | B | D | D |  |
| *Beijerinckia* | PS | B | D | D |  |
| *Chaetomium* | PS | F |  | D |  |
| *Cladosporium* | PS | F |  | D |  |
| *Clonostachys* | PS | F |  | D |  |
| *Colletotrichum* | PS | F |  | D |  |
| *Cryptococcus* | PS | F |  | D |  |
| *Curtobacterium* | PS | B | D |  |  |
| *Enterobacteria* | PS | B |  | D |  |
| *Erwinia* | PS | B | D | D |  |
| *Exiguobacterium* | PS | B | D | D |  |
| *Fusarium* | PS | F |  | D | C |
| *Gluconacetobacter* | PS | B | D | D | Co |
| *Halomonas* | PS | B | D | D |  |
| *Hymenobacter* | PS | B | D | D |  |
| *Hyphomicrobium* | PS | B | D | D | C |
| *Massilia* | PS | B | D |  |  |
| *Methylobacterium* | PS | B | D | D | C |
| *Microbacterium* | PS | B | D | D |  |
| *Mycobacterium* | PS | B | D | D | C |
| *Nocardia* | PS | B | D | D |  |
| *Pseudochrobactrum* | PS | B | D |  |  |
| *Pseudoxanthomonas* | PS | B | D | D | C |
| *Rathayibacter* | PS | B | D |  |  |
| *Rhodopseudomonas* | PS | B | D | D | C |
| *Saccharothrix* | PS | B | D |  |  |
| *Sphingomonas* | PS | B | D | D | C |
| *Stenotrophomonas* | PS | B | D | D | C |
| *Xanthomonas* | PS | B | D | D | C |
| *Acidovorax* | RS | B | D | D | C |
| *Agromyces* | RS | B | D |  |  |
| *Azoarcus* | RS | B | D | D |  |
| *Azospirillum* | RS | B | D | D | C |
| *Bradyrhizobium* | RS | B | D | D | C |
| *Brevibacillus* | RS | B | D | D |  |
| *Brevibacterium* | RS | B | D | D |  |
| *Brevundimonas* | RS | B | D | D | C |
| *Corynebacterium* | RS | B | D | D |  |
| *Cronobacter* | RS | B | D | D |  |
| *Debaryomyces* | RS | F |  | D | C |
| *Dietzia* | RS | B | D |  |  |
| *Escherichia* | RS | B | D | D | C |
| *Halobacillus* | RS | B | D | D |  |
| *Kluyvera* | RS | B | D |  |  |
| *Leptolyngbya* | RS | B |  | D |  |
| *Micrococcus* | RS | B | D | D |  |
| *Neurospora* | RS | F |  | D | C |
| *Nocardioides* | RS | B | D | D |  |
| *Paenibacillus* | RS | B | D | D |  |
| *Rahnella* | RS | B | D | D |  |
| *Ralstonia* | RS | B | D | D | C |
| *Rhizopus* | RS | F |  | D | C |
| *Achromobacter* | PS, RS | B | D | D | C |
| *Acinetobacter* | PS, RS | B | D | D | C |
| *Alcaligenes* | PS, RS | B | D | D |  |
| *Arthrobacter* | PS, RS | B | D | D |  |
| *Bacillus* | PS, RS | B | D | D | C |
| *Burkholderia* | PS, RS | B | D | D | C |
| *Candida* | PS, RS | B |  | D | C |
| *Citrobacter* | PS, RS | B | D | D |  |
| *Enterobacter* | PS, RS | B | D | D | Co |
| *Flavobacterium* | PS, RS | B | D | D | Co |
| *Klebsiella* | PS, RS | B | D | D | C |
| *Pantoea* | PS, RS | B | D | D |  |
| *Penicillium* | PS, RS | F |  | D | C |
| *Pseudomonas* | PS, RS | B | D | D | C |
| *Psychrobacter* | PS, RS | B | D | D |  |
| *Rhizobium* | PS, RS | B | D | D | Co |
| *Rhodococcus* | PS, RS | B | D | D |  |
| *Saccharomyces* | PS, RS | F |  | D | C |
| *Staphylococcus* | PS, RS | B | D | D | C |
| *Streptomyces* | PS, RS | B | D | D | C |
| *Variovorax* | PS, RS | B | D | D | C |

<sup>1</sup>(Bashir et al. 2022; Saeed et al. 2021)

### The Pitcher Metatranscriptome Functions

Our analysis of the pitcher metatranscriptome using the PGPT database revealed major function categories (Table 1, Figure 9). About 50,424 metatranscripts were mapped to PGPT proteins that are associated with different plant growth-promoting functional categories. Globally, 13,519 and 29,028 metatranscripts are mapped to functional groups with direct and indirect effects on plant growth traits, respectively (Figure 9). Among the microbiome traits that have direct effects on plant growth are biofertilization, bioremediation, and phytohormonal plant signals. The microbiome traits that have indirect effect on plant growth include stress control, immune response stimulation, colonizing plant systems and microbial competitive exclusion. As shown in Figure 9, the size of the green circle represents the log scale of the number of metatranscripts assigned to the corresponding function. Altogether, there are 44 major microbiome traits that have direct effect on the host-microbiome and microbiome-microbiome interactions. Traits that are represented by over a thousand metatranscripts include nitrogen acquisition (1003), phosphate solubilization (1751), heavy metals detoxifications (2413), plant vitamin production (1069), neutralizing biotic stress (1578), neutralizing abiotic stress (6230), universal stress response (1062), chemotaxis (1033), plant-derived substrate usage (7067), colonization surface attachment (1485), cell envelop remodeling (1618), biofilm formation (2313), bacterial fitness (2526), and bacterial secretion (1188). These findings reveal the complex functions of the pitcher microbiome at several levels, providing a chance to investigate a wide range of microbiome traits and their host interactions.

**Figure 9:**
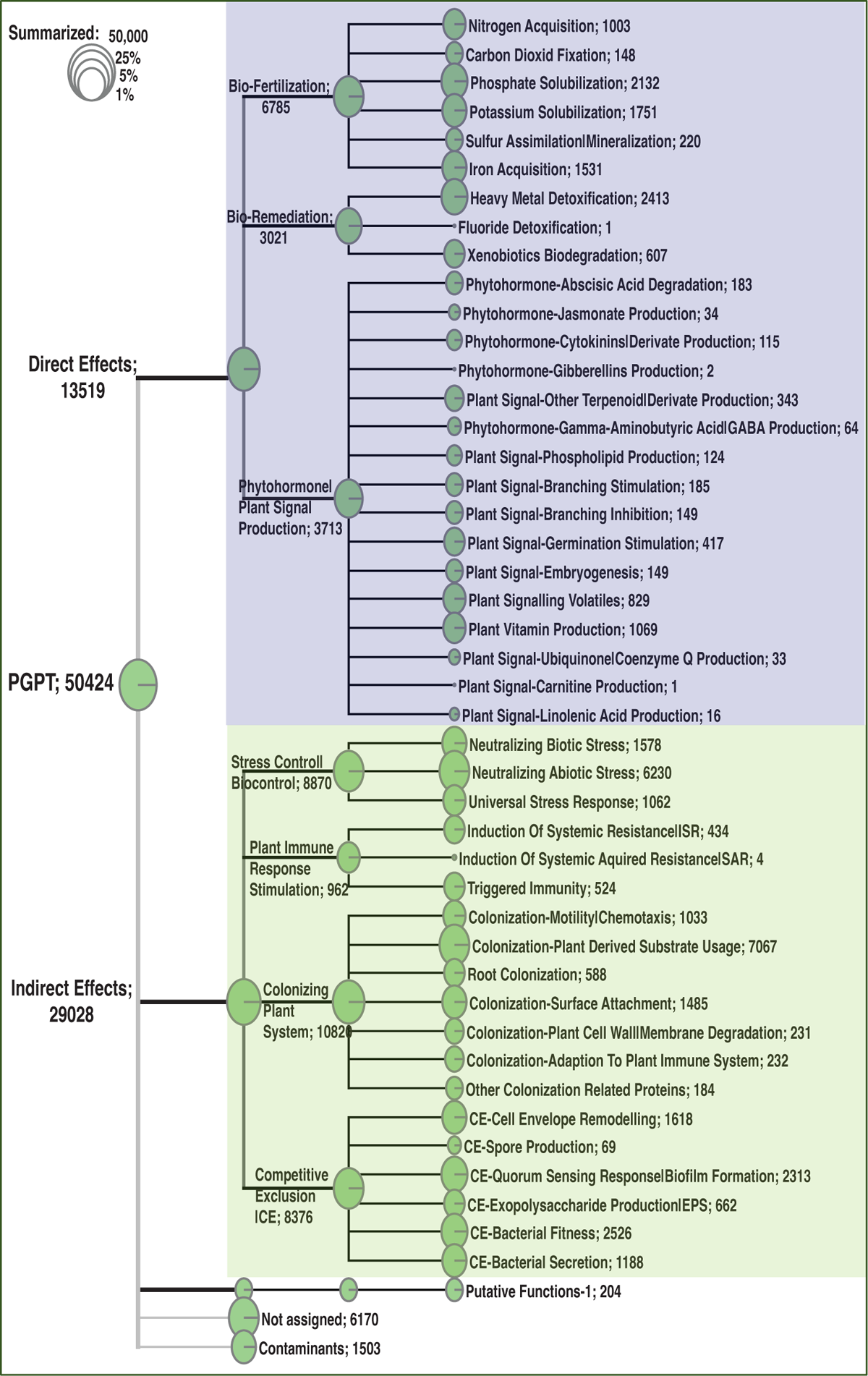
Classification of the major function categories of the pitcher microbiome based on the number of metatranscripts mapped to PGPT ontology. The circles represent the log scale of number of metatranscripts assigned to each function. The number following each function is the number of metatranscripts mapped to this function.

## Discussion

The *Sarracenia* genus relies on its microbiome to facilitate the digestion of trapped prey and to provide important nutrients such as nitrogen (N) for protein synthesis, phosphorus (P) for nucleic acid synthesis, and magnesium (Mg) and iron (Fe) for chlorophyll synthesis. Previous research suggested that factors other than prey capture and colonization by eukaryotic species may affect the recruitment of bacteria to the pitchers (Grothjan and Young 2019). Also, It has been demonstrated that plants have the ability to modify the composition of the microbiomes in their surroundings (Mahnert et al. 2015) and host genetic variations have an impact on the structure and functions of the microbiome in humans (Blekhman et al. 2015; Turnbaugh et al. 2009). Extensive research has been conducted on the convergent evolution of carnivorous plant leaves, particularly those belonging to the *Sarracenia* genus (Wheeler and Carstens 2018; Thorogood et al. 2018; Chomicki et al. 2024). The evolution of the composite trait of the insect-trapping pitchers in carnivorous plants is believed to have occurred through separate modifications of shape, coloration, and biosynthetic pathways (reviewed in (Chomicki et al. 2024)). We are exploring the concept of considering the pitcher microbiome properties as an extension of plant traits that have evolved alongside carnivory as a part of the insect-trapping composite trait. Our hypothesis is that carnivorous plants have gained genetic traits during the convergent evolution of their pitcher trait, enabling them to interact with their vital microbiomes. Our results show that pitcher plants impact the structure and function of their microbiomes, which in turn have a broader effect on their growth, extending beyond mere nutritional support.

### The Host Genotype Significantly Influence the Microbiome Assembly

In their natural habitats, the microbiome diversity diverged among the *Sarracenia* host species (Heil et al. 2022). In this study, we detected statistically significant differences in the structure of the pitcher microbiome communities among the parental species and their F1 and F2 genotypes under the same greenhouse conditions. The finding is supported by the *PERMANOVA* analysis, which indicated that the variations in distribution among genotypes are statistically significant (*p*-value = 0.011). The *PERMDISP* test indicated that there is no significant difference in the dispersion (*p*-value = 0.825) between genotypes, ruling out the possibility that difference in sequencing depth between samples could have produced the observed distribution differences. Both Alpha-diversity and PCoA (Figure 2) analyses unveiled differences in the microbiome diversity among individuals of the same group and between the genotypes. Although the majority of the microbial taxa existed in the microbiomes of all species and genotypes, there were genotype-enriched microbial taxa (Figure 3A, C, and D). The microbiome of the F1 generation differed significantly from all other samples, which could be attributed to its dominant genetics that are expressed in more distinctive traits. The variation in the microbiota between the parental species is smaller than the variation observed between either of them and their F1 and F2 genotypes. This observation is reinforced by the number of distinctive genera, as identified by the LEfSe analysis, between the parental species and their F1 and F2 generation (Figure 3C). The F2 generation exhibits increased genotype complexity and genetic variation, resulting in a wide range of traits, including those that influence the assembly and structure of their microbiome. The abundance patterns of the overall microbiome community hub and connector genera varied significantly among the different genotypes, as shown in Figures 4 and 5. This suggests that the genotypes may influence the development of their respective local community hubs and connectors. By utilizing the *Sarracenia* mapping population, our results suggest that host genetic factors are impacting the observed significant variance in the microbiome structure.

### The Core Microbiomes and Plant Growth-Promoting Traits

The richness and diversity of the sarracenia core microbiome suggests that it possesses a wide range of complex traits and levels of interactions. Over 1900 bacterial genera were identified using 16S the rRNA gene, including 62 ubiquitous genera, 145 genotype-enriched genera, and 58 genera constituting the structural core microbiome. More than 2100 microbial genera were found using the metatranscriptomic data, including 68 bacterial and 22 fungal genera constituting the functional core microbiome. There are 16 genera existed in both structural and functional core microbiomes, including *Acidovorax*, *Bradyrhiobium*, *Bukholderia*, *Caulobacter*, *Chromobacterium*, *Clostridium*, *Cupriavidus*, *Magnetospirilllum*, *Mesorhizobium*, *Novosphingobium*, *Paraburkholderia*, *Pseudomonas*, *Rhizobium*, *Rhodopesudomonas*, *Rickettsia*, and *Stenotrophomonas*. The functional core microbiome relies on the prevalence and abundance of protein-coding metatranscripts, while the identification of the structural microbiome relies on the prevalence and abundance of 16S rRNA gene reads. Many of the core genera especially the prevalent ones are among the group of plant-associated microbes known as plant growth-promoting Rhizobacteria (PGPR) (Beneduzi et al. 2012; Agarwal et al. 2020). As shown in Table 2, we identified the fungal and bacterial taxa in the pitcher microbiome, which have previously been discovered in rhizospheres and/or phyllosphere of other plant species (reviewed in (Bashir et al. 2022; Saeed et al. 2021)). While PGPR has traditionally been found in the rhizosphere, it is not unexpected to discover them in the pitcher microbiome (phyllosphere) due to the fact that this plant species depends on pitcher leaves for acquiring essential nutrients. This finding prompts inquiries into the functional and structural interactions between the rhizosphere and phyllosphere microbiomes while coexisting in the confined microenvironment of the pitchers. The functional traits of these PGPR genera include the ability to enhance plant growth through biofertilization, act as biocontrol agents to combat pests, and serve as biological fungicides to protect their specific host plants.

### Microbiome Community Clusters and Hubs

The network analysis (Figures 4 and 5) unveiled many clusters of microbial communities (File S1), resulting in the identification of community hubs and connector genera. The number of community clusters varied between 3 and 7, depending on the network factors used to identify the clusters. For example, the analysis revealed the presence of five distinct community clusters in the entire network. Additionally, the networks of the top 100 abundant and top 100 participation coefficient nodes (genera) exhibited the identification of six and seven community clusters, respectively. But regardless of the method, there are always distinctive community clusters in the inferred microbiome network. The key characteristic of these communities is the discovery of a total of 27 genera playing the hub and connector roles in these communities. It is crucial to note that the hub and connector genera, based on hubscore, within-model connectivity z-score, and participation coefficient, have a low abundance. This indicates that the genera that perform important roles in the microbiome community are not always the most prevalent genera. Interestingly, the analysis of the normalized abundance of the 27 hub and connector genera in the parents, F1, and F2 genotypes uncovers distinct clusters of genera that vary significantly in their abundance among the genotypes (Figure 5D). This suggests that the host genetic makeup influences the selection of the microbiome hubs and connectors. For instance, the genus *Schlesneria* appeared as a hub genus in the whole network, as well as in the subnetworks, despite not being among the highly abundant genera compared to the other hub and connectors genera (Figure 5D; File S1). As shown in Table 2, the majority of the hub and connector genera are commonly reported as plant-associated genera living in the plant’s phyllosphere and rhizosphere (Mahnert et al. 2018). Overall, the inferred microbiome networks offered insights into the assembly and organization of the pitcher microbiome. This involved the identification of many subcommunities, each with their hub genera, as well as connector genera that facilitate communications between these subcommunities. Interestingly, the community networks also show that the hub and connector genera exhibit varying levels of abundance across different genotypes (Figure 5D), suggesting that the host genotypes may have an impact on their development.

### The Complex Function of Pitcher Microbiome

Although our findings demonstrate a considerable influence of the host on the diversity and organization of the microbiome, we did not discover any notable genotype-related variations in the enriched functional categories in the metatranscriptome (Figure 8). This analysis, however, lacks the level of detail necessary to discover any fine functional differences. Nevertheless, it suggests that even though there may be fine difference in the microbiome functions among genotypes, the overall microbiome traits and functions remain consistent. This could be explained by functional redundancy among the taxa of the pitcher microbiome and a microbiome functional convergence to achieve similar functional traits (Louca et al. 2018, 2016). A similar conclusion was reached by analyzing the microbial communities and functions in *Spu* populations in natural habitats (Grothjan and Young 2019). The functional analysis, however, revealed complex and multifaceted functional traits of the pitcher microbiome (Figure 9). On the host-microbiome interaction level, approximately 13.5% of the metatranscripts that have been classified functionally are related to genes involved in biofertilization activities. These activities encompass nitrogen acquisition, phosphate solubilization, and iron acquisition. The presence of a significant portion of the pitcher microbiome that contributes to plant biofertilization is expected, given that acquiring nutrients from leaves is a key characteristic associated with carnivorous behavior. Nevertheless, the discovery of the pitcher microbiome’s bioremediation (6%) and phytohormonal plant signal generation (7%) activities are novel additions to our understanding. The microbiome has a significant role in the detoxification of heavy metals (2413 metatranscripts) and the degradation of xenobiotics (607 metatranscripts) in the pitcher fluid, which serves to safeguard the host plant. These functions directly impact the fitness and defense of the pitcher plant and can be considered extended plant traits. Approximately 58% of the pitcher metatranscriptome is assigned to a set of functions categorized as traits that have an indirect impact on the host plan (Figure 9). The most prevalent functions, as indicated by the abundance of metatranscripts, are those associated with stress regulation, promotion of plant immune response, and colonization of plant systems. On the microbiome-microbiome interaction level, the analysis revealed that major portions of the metatranscriptome are involved in microbial competitive exclusion activities (8376 metatranscripts) and usage of plant-derived substrates (7067 metatranscripts).

These results offer a comprehensive understanding of the functions of the microbiome in connection to the host plant, microbial community, and environmental conditions inside the pitcher microcosm. Furthermore, they highlight some strategies utilized by the host plant to regulate the assembly of the microbiome, such as plant immunity and the release of extracellular substrates. The pitcher contains a varied microbial population with complicated multiscale functions, which can be used for metagenomic mining to discover new beneficial microbial taxa.

## Conclusion

The variations in the assembly and structure of the microbial communities within the pitchers of the *Sarracenia* mapping population can be attributed, at least in part, to the genetic characteristics of the host. The pitcher microbiome performs multiple functions that have both direct and indirect effects on the host.

Nevertheless, the precise genetic and biochemical pathways via which the host plant interacts with the microbiome are still unidentified. Our goal is to link the extended traits of the *Sarracenia* plant (namely, microbiome traits and functions) to the genetic elements of *Sarracenia* using cross-species QTL linkage mapping. This will enable us to gain insights into the mechanisms underpinning the interactions between the host and its microbiome, which is an essential requirement for the pursuits of microbiome engineering.

## Data availability

The sequencing data will be deposited at NCBI GEO repository and released upon the acceptance of this manuscript.

## Supplementary Materials

### Supplementary Figures

– Figure S1: Metatranscriptome library FA trace.

### Supplementary Files

– File S1: Supplementary File 1 _ microbiome data.
– File S2: Supplementary File 2_metatranscriptome data.

## Reference

Adlassnig W, Peroutka M, Lendi T. 2011. Traps of carnivorous pitcher plants as a habitat: composition of the fluid, biodiversity and mutualistic activities. Annals of Botany 107: 181–194.

Afridi MS, Ali S, Salam A, César Terra W, Hafeez A, Sumaira, Ali B, S. AlTami M, Ameen F, Ercisli S, et al. 2022. Plant Microbiome Engineering: Hopes or Hypes. Biology (Basel) 11: 1782.

Agarwal P, Giri BS, Rani R. 2020. Unravelling the Role of Rhizospheric Plant-Microbe Synergy in Phytoremediation: A Genomic Perspective. Curr Genomics 21: 334–342.

Albert VA, Williams SE, Chase MW. 1992. Carnivorous plants: phylogeny and structural evolution. Science 257: 1491–1495.

Anderson MJ. 2001. A new method for non-parametric multivariate analysis of variance. Austral Ecology 26: 32–46.

Baiser B, Buckley HL, Gotelli NJ, Ellison AM. 2013. Predicting food-web structure with metacommunity models. Oikos 122: 492–506.

Bashir I, War AF, Rafiq I, Reshi ZA, Rashid I, Shouche YS. 2022. Phyllosphere microbiome: Diversity and functions. Microbiological Research 254: 126888.

Bastian M, Heymann S, Jacomy M. 2009. Gephi: an open source socware for exploring and manipulating networks. In Proceedings of the internaAonal AAAI conference on web and social media, Vol. 3 of, pp. 361–362 http://ojs.aaai.org/index.php/ICWSM/arBcle/view/13937 (Accessed December 18, 2023).

Beneduzi A, Ambrosini A, Passaglia LMP. 2012. Plant growth-promoting rhizobacteria (PGPR): Their potential as antagonists and biocontrol agents. Genet Mol Biol 35: 1044–1051.

Bieleston LS, Wolock CJ, Yahya BE, Chan XY, Chan KG, Pierce NE, Pringle A. Convergence between the microcosms of Southeast Asian and North American pitcher plants. eLife 7: e36741.

Blekhman R, Goodrich JK, Huang K, Sun Q, Bukowski R, Bell JT, Spector TD, Keinan A, Ley RE, Gevers D, et al. 2015. Host genetic variation impacts microbiome composition across human body sites. Genome Biology 16: 191.

Boyer T, Carter R. 2011. Community Analysis of Green Pitcher Plant (Sarracenia oreophila) Bogs in Alabama. Castanea 76: 364–376.

Bray JR, Curtis JT. 1957. An Ordination of the Upland Forest Communities of Southern Wisconsin. Ecological Monographs 27: 325–349.

Bushmanova E, Antipov D, Lapidus A, Prjibelski AD. 2019. rnaSPAdes: a de novo transcriptome assembler and its application to RNA-Seq data. GigaScience 8: giz100.

Chao A. 1984. Nonparametric Estimation of the Number of Classes in a Population. Scandinavian Journal of StaAsAcs 11: 265–270.

Chomicki G, Burin G, Busta L, Gozdzik J, Jeeer R, Mortimer B, Bauer U. 2024. Convergence in carnivorous pitcher plants reveals a mechanism for composite trait evolution. Science 383: 108–113.

Clauset A, Newman MEJ, Moore C. 2004. Finding community structure in very large networks. Phys Rev E 70: 066111.

Dobin A, Davis CA, Schlesinger F, Drenkow J, Zaleski C, Jha S, Batut P, Chaisson M, Gingeras TR. 2013. STAR: ultrafast universal RNA-seq aligner. BioinformaAcs 29: 15–21.

Ellison AM, Buckley HL, Miller TE, Gotelli NJ. 2004. Morphological variation in *Sarracenia purpurea* (Sarraceniaceae): geographic, environmental, and taxonomic correlates. Am J Bot 91: 1930–1935.

Ellison AM, Gotelli NJ, Błędzki LA, Butler JL. 2021. Regulation by the Pitcher Plant Sarracenia purpurea of the Structure of its Inquiline Food Web. amid 186: 1–15.

Ellison AM, Gotelli NJ, Brewer JS, Cochran-Stafira DL, Kneitel JM, Miller TE, Worley AC, Zamora R. 2003a. The evolutionary ecology of carnivorous plants. In Advances in Ecological Research, Vol. 33 of, pp. 1–74, Academic Press https://www.sciencedirect.com/science/arBcle/pii/S0065250403330090 (Accessed March 16, 2024).

Ellison AM, Gotelli NJ, Brewer JS, Cochran-Stafira DL, Kneitel JM, Miller TE, Worley AC, Zamora R. 2003b. The evolutionary ecology of carnivorous plants. Advances in Ecological Research*, Vol* 33 **33**: 1–74.

Fisher RA. 1936. THE USE OF MULTIPLE MEASUREMENTS IN TAXONOMIC PROBLEMS. Annals of Eugenics 7: 179–188.

Fu C-N, Wicke S, Zhu A-D, Li D-Z, Gao L-M. 2023. Distinctive plastome evolution in carnivorous angiosperms. BMC Plant Biology 23: 660.

Furches MS, Small RL, Furches A. 2013. Hybridization leads to interspecific gene flow in Sarracenia (Sarraceniaceae). Am J Bot 100: 2085–2091.

Gautam A, Zeng W, Huson DH. 2023. DIAMOND + MEGAN Microbiome Analysis. In Metagenomic Data Analysis (ed. S. Mitra), pp. 107–131, Springer US, New York, NY 10.1007/978-1-0716-3072-3_6 (Accessed May 6, 2024).

Ge SX, Jung D, Yao R. 2020. ShinyGO: a graphical gene-set enrichment tool for animals and plants. BioinformaAcs 36: 2628–2629.

Gower JC. 1966. Some distance properties of latent root and vector methods used in multivariate analysis. Biometrika 53: 325–338.

Grothjan JJ, Young EB. 2022. Bacterial Recruitment to Carnivorous Pitcher Plant Communities: Identifying Sources Influencing Plant Microbiome Composition and Function. Front Microbiol 13: 791079.

Grothjan JJ, Young EB. 2019. Diverse microbial communities hosted by the model carnivorous pitcher plant Sarracenia purpurea: analysis of both bacterial and eukaryotic composition across distinct host plant populations. PeerJ 7: e6392.

Guimera R, Nunes Amaral LA. 2005. Functional cartography of complex metabolic networks. nature 433: 895–900.

Heil JA, Wolock CJ, Pierce NE, Pringle A, Bieleston LS. 2022. Sarracenia pitcher plant-associated microbial communities differ primarily by host species across a longitudinal gradient. Environmental Microbiology 24: 3500–3516.

Ke J, Wang B, Yoshikuni Y. 2021. Microbiome Engineering: Synthetic Biology of Plant-Associated Microbiomes in Sustainable Agriculture. Trends in Biotechnology 39: 244–261.

Kers JG, Saccenti E. 2022. The Power of Microbiome Studies: Some Considerations on Which Alpha and Beta Metrics to Use and How to Report Results. Front Microbiol 12: 796025.

Kim D, Song L, Breitwieser FP, Salzberg SL. 2016. Centrifuge: rapid and sensitive classification of metagenomic sequences. Genome Res. https://genome.cshlp.org/content/early/2016/11/16/gr.210641.116 (Accessed May 10, 2022).

Koopman MM, Carstens BC. 2011. The Microbial Phyllogeography of the Carnivorous Plant Sarracenia alata. Microbial Ecology 61: 750–758.

Koopman MM, Fuselier DM, Hird S, Carstens BC. 2010. The Carnivorous Pale Pitcher Plant Harbors Diverse, Distinct, and Time-Dependent Bacterial Communities. Appl Environ Microbiol 76: 1851–1860.

Kopylova E, Noé L, Touzet H. 2012. SortMeRNA: fast and accurate filtering of ribosomal RNAs in metatranscriptomic data. BioinformaAcs 28: 3211–3217.

Kruskal WH, Wallis WA. 1952. Use of Ranks in One-Criterion Variance Analysis. Journal of the American StaAsAcal AssociaAon 47: 583–621.

Lahti L, Sheey S. 2018. Introduction to the microbiome R package. *Preprint at* https://microbiomegithubio/tutorials. https://s3.jcloud.sjtu.edu.cn/899a892efef34b1b944a19981040f55b-oss01/bioconductor/3.10/bioc/vigneees/microbiome/inst/doc/vigneee.html (Accessed December 18, 2023).

Louca S, Jacques SMS, Pires APF, Leal JS, Srivastava DS, Parfrey LW, Farjalla VF, Doebeli M. 2016. High taxonomic variability despite stable functional structure across microbial communities. Nat Ecol Evol 1: 1–12.

Louca S, Polz MF, Mazel F, Albright MBN, Huber JA, O’Connor MI, Ackermann M, Hahn AS, Srivastava DS, Crowe SA, et al. 2018. Function and functional redundancy in microbial systems. Nat Ecol Evol 2: 936–943.

Mahnert A, Haratani M, Schmuck M, Berg G. 2018. Enriching Beneficial Microbial Diversity of Indoor Plants and Their Surrounding Built Environment With Biostimulants. Front Microbiol 9: 2985.

Mahnert A, Moissl-Eichinger C, Berg G. 2015. Microbiome interplay: plants alter microbial abundance and diversity within the built environment. Front Microbiol 6. https://www.fronBersin.org/journals/microbiology/arBcles/10.3389/fmicb.2015.00887/full (Accessed May 18, 2024).

Malmberg RL, Rogers WL, Alabady MS. 2018. A carnivorous plant genetic map: pitcher/insect-capture QTL on a genetic linkage map of Sarracenia. Life Sci Alliance 1. http://www.life-science-alliance.org/content/1/6/e201800146.abstract.

McMurdie PJ, Holmes S. 2013. phyloseq: an R package for reproducible interactive analysis and graphics of microbiome census data. PloS one 8: e61217.

Morales Moreira ZP, Chen MY, Yanez Ortuno DL, Haney CH. 2023. Engineering plant microbiomes by integrating eco-evolutionary principles into current strategies. Current Opinion in Plant Biology 71: 102316.

Munch K, Boomsma W, Willerslev E, Nielsen R. 2008. Fast phylogenetic DNA barcoding. Phil Trans R Soc B 363: 3997–4002.

ONT 16S workflow 16S Workflow. 2019. EPI2ME WIMP workflow: quantitative, real-Bme species identification from metagenomic samples. Oxford Nanopore Technologies. https://nanoporetech.com/resource-centre/epi2me-wimp-workflow-quanBtaBve-real-Bme-species-identificaBon-metagenomic (Accessed February 23, 2024).

Patz S, Gautam A, Becker M, Ruppel S, Rodríguez-Palenzuela P, Huson D. 2021. PLaBAse: A comprehensive web resource for analyzing the plant growth-promoting potential of plant-associated bacteria. 2021.12.13.472471. https://www.biorxiv.org/content/10.1101/2021.12.13.472471v1 (Accessed May 6, 2024).

Patz S, Rauh M, Gautam A, Huson DH. 2024. mgPGPT: Metagenomic analysis of plant growth-promoting traits. 2024.02.17.580828. https://www.biorxiv.org/content/10.1101/2024.02.17.580828v1 (Accessed May 6, 2024).

Saeed Q, Xiukang W, Haider FU, Kučerik J, Mumtaz MZ, Holatko J, Naseem M, Kintl A, Ejaz M, Naveed M, et al. 2021. Rhizosphere Bacteria in Plant Growth Promotion, Biocontrol, and Bioremediation of Contaminated Sites: A Comprehensive Review of Effects and Mechanisms. Int J Mol Sci 22: 10529.

Segata N, Izard J, Waldron L, Gevers D, Miropolsky L, Garree WS, Hueenhower C. 2011. Metagenomic biomarker discovery and explanation. Genome Biol 12: R60.

Shannon CE. 2001. A mathematical theory of communication. SIGMOBILE Mob Comput Commun Rev 5: 3–55.

Thorogood CJ, Bauer U, Hiscock SJ. 2018. Convergent and divergent evolution in carnivorous pitcher plant traps. New Phytologist 217: 1035–1041.

Trivedi P, Leach JE, Tringe SG, Sa T, Singh BK. 2020. Plant-microbiome interactions: from community assembly to plant health. Nat Rev Microbiol 18: 607–621.

Turnbaugh PJ, Hamady M, Yatsunenko T, Cantarel BL, Duncan A, Ley RE, Sogin ML, Jones WJ, Roe BA, Affourtit JP, et al. 2009. A core gut microbiome in obese and lean twins. Nature 457: 480–484.

Volvenko IV. 2014. General Paeerns of Spatial Distribution of the Integral Characteristics of Benthic Macrofauna of the Northwestern Pacific and Biological Structure of Ocean. Open Journal of Ecology 4: 196–213.

Wen T, Xie P, Yang S, Niu G, Liu X, Ding Z, Xue C, Liu Y-X, Shen Q, Yuan J. 2022. ggClusterNet: An R package for microbiome network analysis and modularity-based multiple network layouts. iMeta 1: e32.

Wheeler GL, Carstens BC. 2018. Evaluating the adaptive evolutionary convergence of carnivorous plant taxa through functional genomics. PeerJ 6: e4322.

Whitham TG, Bailey JK, Schweitzer JA, Shuster SM, Bangert RK, LeRoy CJ, Lonsdorf EV, Allan GJ, DiFazio SP, Poes BM, et al. 2006. A framework for community and ecosystem genetics: from genes to ecosystems. Nature Reviews GeneAcs 7: 510.

Whitham TG, DiFazio SP, Schweitzer JA, Shuster SM, Allan GJ, Bailey JK, Woolbright SA. 2008. Extending Genomics to Natural Communities and Ecosystems. Science 320: 492–495.

Wilcoxon F. 1992. Individual Comparisons by Ranking Methods. In Breakthroughs in StaAsAcs (eds. S. Kotz and N.L. Johnson), Springer Series in StaAsAcs, pp. 196–202, Springer New York, New York, NY http://link.springer.com/10.1007/978-1-4612-4380-9_16 (Accessed December 18, 2023).

Yan L. 2021. ggvenn: Draw Venn Diagram by ‘ggplot2.’ R Package Version 19.

